# PQBP1-dependent alternative RNA splicing underlies high calorie diet-induced cognitive impairment

**DOI:** 10.1101/2025.06.02.657545

**Authors:** Yong Huang, Hidenori Homma, Xigui Chen, Hikari Tanaka, Kyota Fujita, Albert R. La Spada, Hitoshi Okazawa

## Abstract

High calorie-high fat diet (HFD) has been implicated as a pathological modifier of brain diseases including neurodegenerative dementias, but the detailed molecular mechanisms remain largely unknown. Here we report that HFD suppresses PPARγ-mediated transcriptional expression of *PQBP1*, a RNA splicing factor implicated in human intellectual disability and Alzheimer’s disease. RNAseq-based comprehensive analyses of alternative RNA splicing (AS) in HFD-fed mice for 1 or 6 weeks and in PQBP1-cKO mice reveal their common changes, which weigh on synapse-related genes. Betweenness-based extraction of core molecules from the common changes reveals CASK, Cacnb1 and Cyfip2 as key molecules of the network. Both CASK and Cacnb1 regulate STXBP1, a causative gene for infantile epilepsy syndrome and an essential factor for synapse vesicle release, via their direct interaction. In addition, our analysis suggests that Syt1 plays a role specifically in HFD for 1 week. HFD-induced AS isoforms of CASK, Cacnb1, Cyfip2 and Syt1 impair pre-synapse vesicle release in primary neurons. AAV-PQBP1, AAV-CASK, AAV-Cacnb1, AAV-Cyfip2 or AAV-Syt1 rescues synapse and/or cognitive dysfunctions in HFD mice, genetically supporting the pathological PQBP1-presynase axis in HFD. Moreover, immunohistochemistry experiments suggest that the pathological axis plays roles not only in excitatory neurons, but also in inhibitory neurons of the brain. Collectively, our results unravel a novel molecular mechanism for brain dysfunction when mice are exposed to a HFD.

## Introduction

HFD has been implicated in impairment of brain cognitive functions, and various factors in response to HFD are thought to affect brain function indirectly^1,2^. For instance, HFD decreases synapse marker proteins based upon immunohistochemistry^3–5^, induces insulin resistance^6,7^, activates microglia^4^, induces neuroinflammation^8–10^, alters synaptic transmission in physiological analysis^11,12^, and impairs cognitive function^4,7^. Meanwhile, direct molecular mechanisms connecting HFD to synapse dysfunction in neurons and finally linking HFD to cognitive impairment remain totally unknown. In parallel, HFD has been implicated in exacerbation of senile dementia, such as Alzheimer’s disease (AD), basically through analogously indirect mechanisms^13–17^ ApoE4, the major genetic risk factor that increases the incidence of AD to 2- or 3-fold in a hemizygous dose^18,19^, is another clue for the significance of lipid metabolism for cognitive function^20^. However, evidence accumulated thus far is mostly linked to metabolism of amyloid beta (Aβ) or Tau; hence, the role of Aβ-independent mechanisms remains elusive^18,21^. In other words, it is not known whether Aβ or Tau has a similar molecular target to that of HFD.

PQBP1 was discovered as a binding protein to the polyglutamine (polyQ) tract domain shared by various polyQ disease-causative genes and some transcription-related factors^22,23^ and later identified as a transcription-splicing coupling factor^24–26^. PQBP1 influences RNA splicing patterns transcribed from multiple genes, and such alternative splicing (AS) targets of PQBP1 include cell cycle genes in neural stem cells^27^, a regulatory factor involved in nonsense mRNA mediated decay (NMD)^28^, neurite outgrowth-related genes in primary neurons^29^, synapse-related genes in cortical neurons^30^, and Bcl-2-associated X protein (BAX) in ovarian cancer cells^31^. Synapse dysfunction is caused by abnormal AS of such synapse-related genes due to deficiency of PQBP1 and of its binding partner SRRM2 in intracellular amyloid-positive neurons of human AD patients as well as iPSC-derived neurons from human AD patients^30^. Meanwhile, PQBP1 has a different role as an intracellular receptor for cDNA and protein of HIV1 in innate immune cells^32,33^, and also for tau protein of neurodegenerative diseases in brain microglia^34^. Moreover, PQBP1 is involved in translational elongation^35^ and stress granule homeostasis^36^, but its transcriptional regulation has not been fully investigated.

Recent studies suggest that an imbalance of excitatory and inhibitory neuronal activities (E-I imbalance) is a critical factor affecting cognitive function and protein aggregation pathology in AD^38–40^, and the relative decrease of inhibitory neuron activity exacerbates AD^41–43^. Micro-neural circuits composed of excitatory neurons and inhibitory neurons regulate context-dependent excitatory activity, and the abnormal state could lead to hyperexcitability, loss of gamma oscillations, and structural degradation of GABAergic synapses^44^. In regard to HFD-related mechanisms, hyperphagia is related to functional and/or anatomical deficiencies in specific neurons such as PNOCARC or GABAergic inhibitory neurons^45,46^. However, molecular mechanisms in the opposite direction, i.e. from high calorie-high fat feeding to the E-I imbalance or functional deficiency of inhibitory neurons, are remain ill-defined.

E-I imbalance and functional deficiency of inhibitory neurons are also suggested in intellectual disability (ID) and autism spectrum disorder, including Fragile X syndrome (FXS)^47–50^. Fragile X mental retardation protein (FMRP), the causative gene for FXS, is expressed in GABAergic inhibitory neurons, and FMRP deficiency in full mutation patients may impair GABAergic neurotransmission^51,52^. Meanwhile, the other genes of intellectual disability and autism spectrum disorder, including PQBP1, have not been sufficiently addressed from the aspect of E-I balance or inhibitory neurons. Intriguingly, PQBP1 is decreased in human cortical neurons of AD patients, and human neurons derived from induced pluripotent stem cells (iPSCs) carrying genome-edited AD mutations^30^ and PQBP1 gene mutations cause lean body in human ID patients^53^ and abnormal lipid metabolism in nematode models^54^, suggesting that PQBP1 might be a missing link molecule between metabolism and E-I imbalance.

In this study, starting from our discovery that HFD suppresses PPARγ-mediated transcription of PQBP1 gene, NGS-RNAseq-based AS analysis elucidated influences of PQBP1 on a core molecular network related to synapse function, and mathematical analysis of this molecular network revealed three key genes: Calcium/Calmodulin Dependent Serine Protein Kinase (Cask), cytoplasmic FMR1-interacting protein 2 (*Cyfip2*), calcium voltage-gated channel auxiliary subunit beta 1 (*Cacnb1*), and a common binding target, Syntaxin Binding Protein 1 (*STXBP1*), based on their shared protein structures. STXBP1 is a factor known to regulate synapse vesicle release via Syntaxin1 in inhibitory neurons^55,56^. Interestingly, *Cyfip2* and *STXBP2* are causative genes for infantile epilepsies such as West syndrome^57^ and of developmental and epileptic encephalopathy 4 or Ohtahara syndrome^58–61^ respectively. In addition, our analysis suggests that synaptotagmin 1 (*Syt1*) also plays a short term role lasting about one week in the HFD response. To confirm the physiological effects of these targets and pathway, we performed AAV-mediated rescue experiments of HFD mice and documented that PQBP1 or its immediate downstream target proteins can reverse cognitive dysfunction. Collectively, our results indicate that the PQBP1-Cacnb1/CASK-STXBP1 axis plays a critical role in HFD-induced cognitive dysfunction.

## Results

### HFD suppresses expression of PQBP1 in neurons

One of the phenotypes of human patients with *PQBP1* gene mutations is lean body mass^53,62^. In addition, we previously found that nematode models with mutations of the nematode Pqbp1 homologue gene show abnormalities in lipid storage^54^. These results prompted us to investigate the relationship between PQBP1 and high calorie-high fat diet (HFD). After confirming that HFD induced body weight gain, hyper-glycemia, and hyper-insulinemia (Supplementary Figure 1), we first examined how HFD affects gene and protein expression of PQBP1, and our western blot revealed that pretreatment of HFD for 6 weeks (HFD-6W) and even for 1 week (HFD-1W) (Figure 1a) suppressed protein levels of PQBP1 in the cerebral cortex of C57BL/6 wild-type mice at 12 weeks of age (Figure 1b). Reverse transcription polymerase chain reaction (RT-PCR) analysis revealed reduced *PQBP1* mRNA levels (Figure 1c), suggesting that the protein reduction was mostly likely due to transcription dysregulation of the *PQBP1* gene or post-transcriptional decreased stability of the *PQBP1* mRNA.

**Figure 1:**
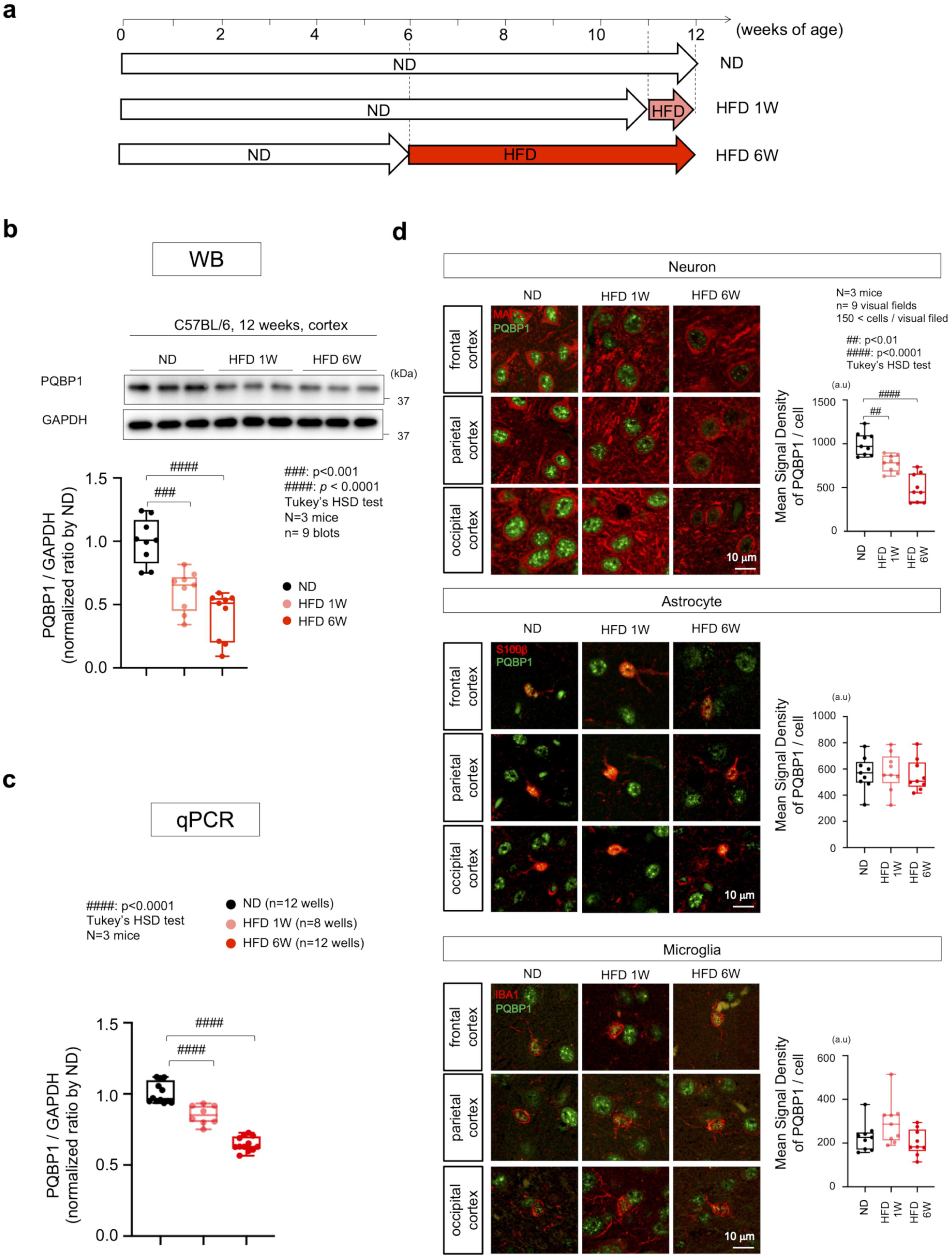
High calorie diet suppresses protein and gene expressions of PQBP1. a) Protocol for feeding after lactation period. ND: mice fed with normal diet from 4 to 12 weeks of age. HFD-1W: mice fed with normal diet from 4 to 11 weeks and with HFD from 11 to 12 weeks of age. HFD-6W: mice fed with normal diet from 4 to 6 weeks and with HFD from 6 to 12 weeks of agesaralab) Western blot analysis of PQBP1 protein with total cerebral cortex tissues from ND, HFD-1W and HFD-6W mouse groups.saralac) RT-qPCR data of *PQBP1* mRNA with total cerebral cortex tissues of ND, HFD-1W and HFD-6W mouse groups.saralad) Immunohistochemistry of PQBP1 in neurons, astrocytes and microglia in frontal cortex. Right graphs show mean signal intensity per cell, in which signal/cell was subtracted by background signal and the mean value was calculated in a visual field. 9 visual filed were examined in frontal cortex of three mice.

We further performed immunohistochemistry to examine which type of cells in the brain were affected by HFD (Figure 1d). Both in HFD-1W and HFD-6W, signal intensities of PQBP1 protein were decreased selectively in cortical neurons. PQBP1-mediated Tau-recognition and innate immune response in brain microglia are attracting attention recently^34^, but the protein levels were not changed in microglia or astrocytes (Figure 1d). Therefore, we focused on neuronal regulation of PQBP1 and the consequences to neurons.

### PPAR**γ** mainly mediates HFD-induced PQBP1 reduction

Next, we addressed the mechanism by which HFD suppressed PQBP1 trascript expression. Several software programs exist to find transcription factor binding sites (e.g. TFBIND: https://tfbind.hgc.jp/, rVista: https://rvista.dcode.org/), and these algorithms revealed a typical consensus sequence for peroxisome proliferator-activated receptors (PPARs)^63,64^ in the promoter upstream of the *PQBP1* gene (Figure 2a). PPAR was originally found as an orphan receptor for diverse hepatocarcinogens^65^, and the family members turned out to be central regulators of body metabolism via adipocyte, acyl-CoA oxidase, or beta-oxidation pathways^66–68^. It is known that the MEK-ERK pathway is activated in HFD and promotes PPARγ phosphorylation at Ser273 to regulate PPARγ function to suppress insulin sensitivity^69^. Each PPAR (PPARα, PPARδ and PPARγ) can form a hetrodimer with the retinoid X receptors (RXRs)^67^ which cooperate to regulate gene expression as a transcription factor complex^70^. Therefore, we hypothesized that PPARγ might enhance transcription of PQBP1 gene and that PPARγ phosphorylation at Ser273 might suppress the transcriptional activity.

**Figure 2:**
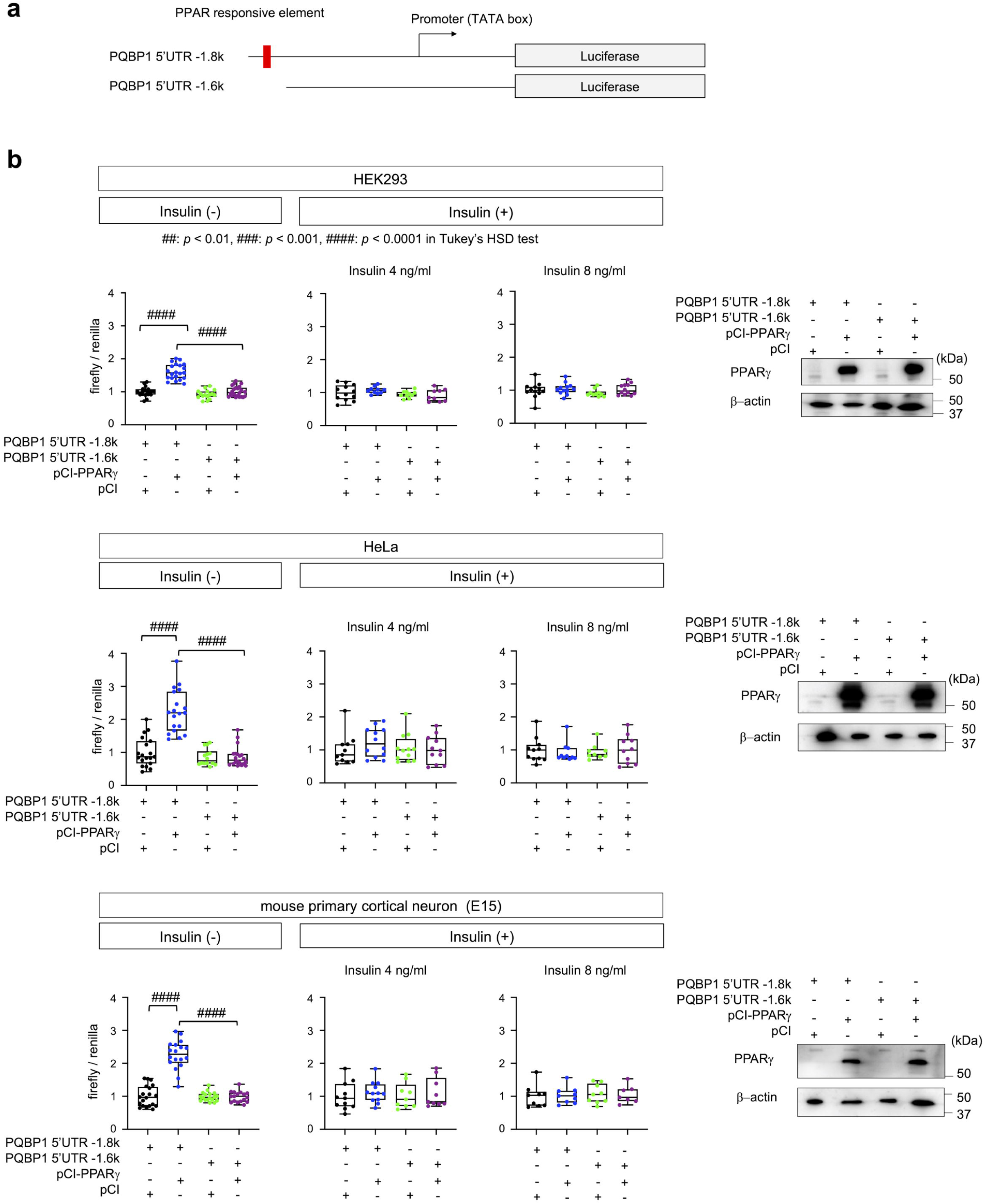
PPARγ regulates PQBP1 gene expression. a) Reporter vectors possessing 1.8 kb or 1.6 kb of the upstream region of mouse *PQBP1* gene were used for luciferase assay. The PPAR-responsive element (<u>AGGTCA</u>G<u>AGGTCA</u>) completely matching with the consensus sequence (<u>AGGTCA</u>N<u>AGGTCA</u>) ^63,64^ is shown with a red box in the scheme.saralab) Luciferase assay was performed in HEK293 cells, HeLa cells and mouse primary cortical neurons, to examine the effect of PPARγ on *PQBP1* gene expression. Right panels show expression of PPAR in the absence of insulin.

To examine this hypothesis, we co-transfected an expression vector of PPARγ together with a luciferase reporter construct containing the upstream promoter region of the PQBP1 gene with or without the consensus PPARγ binding sequence (Figure 2b). In addition to primary cortical neurons prepared from mouse E15 embryos, we performed the luciferase assay in HEK293 cells and HeLa cells as the viability of primary cortical neurons might affect the result of the luciferase assay (Figure 2b). We also added insulin to culture medium to activate the MEK/ERK pathway that phosphorylates PPARγ at Ser273 to influence the transcription factor complex activity. As expected, in all three types of cells, PPARγ activated transcription of PQBP1, transactivation by PPARγ was mediated by the consensus sequence for PPARγ binding, and insulin abolished PPARγ transactivation (Figure 2b).

We performed a similar assay with expression vectors for the other PPAR family transcription factors (Supplementary Figure 2). Though endogenous expression levels of PPARγ, PPARα, and PPARδ are different according to the database (Supplementary Figure 2a), as expected, PPARα and PPARδ similarly enhanced transcription from PQBP1 gene (Supplementary Figure 2b, c). However, the extent of transactivation by PPARα and PPARδ was much less than that of PPARγ in our experimental conditions (Supplementary Figure 2 v.s. Figure 2).

In addition, we performed the luciferase assay of PPARγ in primary mouse microglia (Supplementary Figure 3). As expected from the immunohistochemistry analysis of PQBP1 in the brain microglia of HFD-6W mice (Figure 1d), we did not detect transcription activation of PQBP1 promoter/enhancer by PPARγ (Supplementary Figure 3).

### HFD affects dendritic spines of excitatory synapses

Because PQBP1 regulates splicing of synapse-related genes^27,30^, we suspected that the HFD-induced decrease of PQBP1 protein in neurons may lead to morphological and functional impairments of neuronal synapses. As expected, two-photon microscopy of dendritic spines visualized by AAV1-EGFP revealed a decrease in the number of dendritic spines in the first layer of parietal cerebral cortex (Figure 3a), though morphological features of a single spine did not change (Figure 3a). AAV2-VAMP2-mCherry injected simultaneously also revealed the decrease of EGFP-mCherry-double-positive mature synapses (Figure 3a). Live imaging of the same dendritic spines by two-photon microscopy revealed an increase of spine elimination, which likely explains the decrease of dendritic spines in HFD-1W and HFD-6W mice (Figure 3b). The decrease of excitatory synapses in HFD-1W and HFD-6W mice was further confirmed by western blot analysis of PSD95 (Figure 3c), a marker protein of the post-synapse density (PSD) that forms an excitatory synapse with the pre-synaptic terminal. In consistence, western blot also revealed reduction of VAMP2, a marker protein of the pre-synaptic terminal, in HFD-1W and HFD-6W mice (Figure 3c).

**Figure 3:**
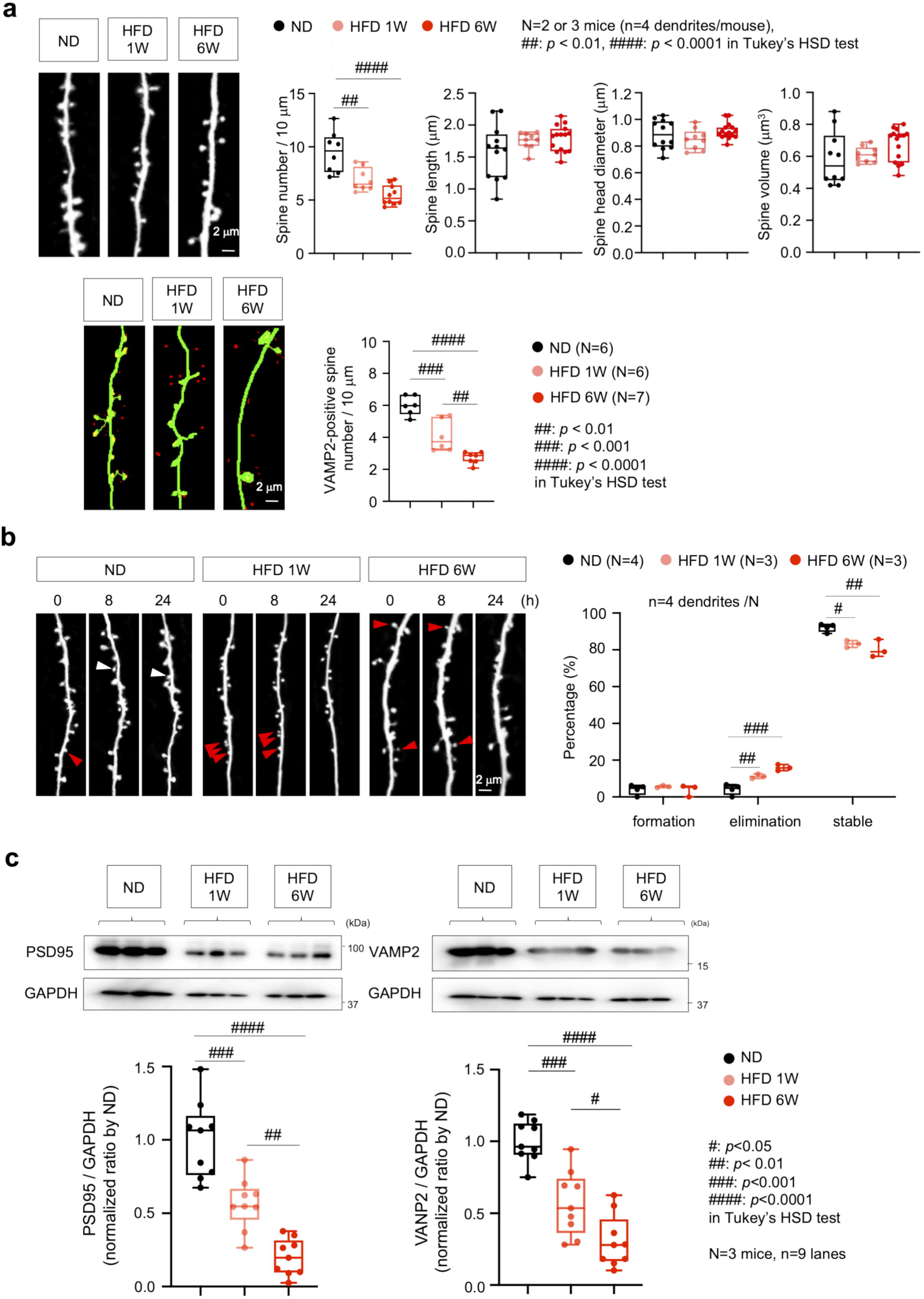
High calorie diet impairs dendritic spines. a) Upper panels show density, length, diameter and volume of dendritic spines in the mouse splenial cortex. The density was changed while the other parameters were not significantly affected by HFD. Lower panels show VAMP2-positive dendritic spines of mice co-injected by AAV2-VAMP2-mCherry and AAV1-EGFP directed by synapsin-1 promoter.saralab) Dynamics of dendritic spines during 24 hours of observation by two-photon microscopy. Eliminated spines during 24 hours are indicated with red arrowheads. Elimination and stability was changed while formation was not affected by HFD.saralac) Western blot of cerebral cortex with a post-synapse marker PSD95. Three blots were independently performed with a mouse. Right panels show quantitative analysis of signal intensities of the PSD95 bands obtained from 9 blots of 3 mice.saralad) Re-examination of exon skipping by an independent group of Syn1-Cre/PQBP1-cKO mice (N=3), HFD-1W mice (N=3), HFD-6W mice (N=3) and C57BL/6J mice (N=3). Odds ratios of each mice vs C57BL/6J mice are shown with p-value by Fisher’s exact test and FDR by Benjamini-Hochberg procedure.

### Target genes of HFD-induced impairment of alternative RNA splicing

To unravel which molecules were affected in RNA splicing under HFD-induced PQBP1-deficeincy to cause such synapse impairments, we performed RNAseq analysis of AS^30^ in order to map exon skipping (Figure 4a). To do so, we analyzed cerebral cortex samples prepared from mice with HFD for 1 or 6 weeks (HFD-1W or HFD-6W) together with mice with cortical neuron-specific depletion of PQBP1 (synapsin 1-Cre mediated conditional knockout mouse of PQBP1, Syn-cKO), and we extracted the common AS profiles that were shared between HFD mice and PQBP1-deficient mice (Figure 4b). Consequently, 716 genes in Syn-cKO and HFD-1W mice displayed shared RNA splicing alterations, and 695 genes in Syn-cKO and HFD-6W mice displayed shared RNA splicing alterations. These genes were identified and used in GO-based cluster analysis (Figure 4b, Supplementary Figure 4, 5). In both groups of commonly changed genes, GO-based cluster analysis generated a group of 162 and 150 genes repectively enriched for “synapse-related signals” (Figure 4b, Supplementary Figure 4, 5). Genes involved in “synapse-related signals” were then subjected to molecular network analysis on protein-protein interaction (PPI) databases (integrated database of the Genome Network Project (GNP: https://cell-innovation.nig.ac.jp/GNP/index_e.html), which includes BIND (http://www.bind.ca/), BioGrid (http://www.thebiogrid.org/), HPRD (http://www.hprd.org/), IntAct (http://www.ebi.ac.uk/intact/site/index.jsf), and MINT (https://mint.bio.uniroma2.it/)), and their expected functional significances were ranked by a “betweenness score”, a parameter reflecting impact on the PPI network (Figure 4b). Moreover, we selected 10 and 15 genes (top 10%) based upon betweenness scores from the comparison of Syn-cKO and HFD-1W and that of Syn-cKO and HFD-6W (Figure 4b, c). Surprisingly, 6 genes were shared in common of these 10 and 15 genes (Figure 4b, c). Furthermore, the top three genes (*Cask*, *Cyfip2* and *Cacnb1*) were closely related in the PPI network predicted by String (https://string-db.org/) (Supplementary Figure 6a), which included amyloid precursor protein (APP) interestingly (Supplementary Figure 6a). Cask and Cacnb1 were connected via STXBP1 in the PPI network (Supplementary Figure 6a) that was also selected as one of the final 6 genes (Figure 4c). Cask and Cacnb1 are membrane-associated guanylate kinase (MAGUK) family members possessing a Guanylate kinase-like (GuK) domain and SH3 protein binding domains (Supplementary Figure 7). The skipping of exon 13 in Cask leads to loss of the L27 domain and a drastic change in protein conformation according to the prediction by Alphafold v2.2.4 (Supplementary Figure 7). The skipping of exon 7 in Cacnb1 also leads to loss of an IDP domain and a drastic change in protein conformation according to Alphafold v2.2.4 (Supplementary Figure 7). Consistently, co-immunoprecipitation of His-STXBP1 with Myc-Cacnb1 or with Myc-Cask revealed interaction of STXBP1 with full-length forms, but not with the exon skipped Cask or Cacnb1 isoforms (Supplementary Figure 8). We further examined the three candidate genes of PQBP1 downstream with independent mouse groups of Syn-cKO, HFD-1W, and HFD-6W, and observed reproducible results of CASK and partially reproducible results of Cyfip2 and Cacnb1 (Figure 4d). Collectively these results suggested a molecular network that was distributed around the most reliable molecule Cask as the key target gene of HFD-induced and PQBP1-mediated impairment of RNA splicing.

**Figure 4:**
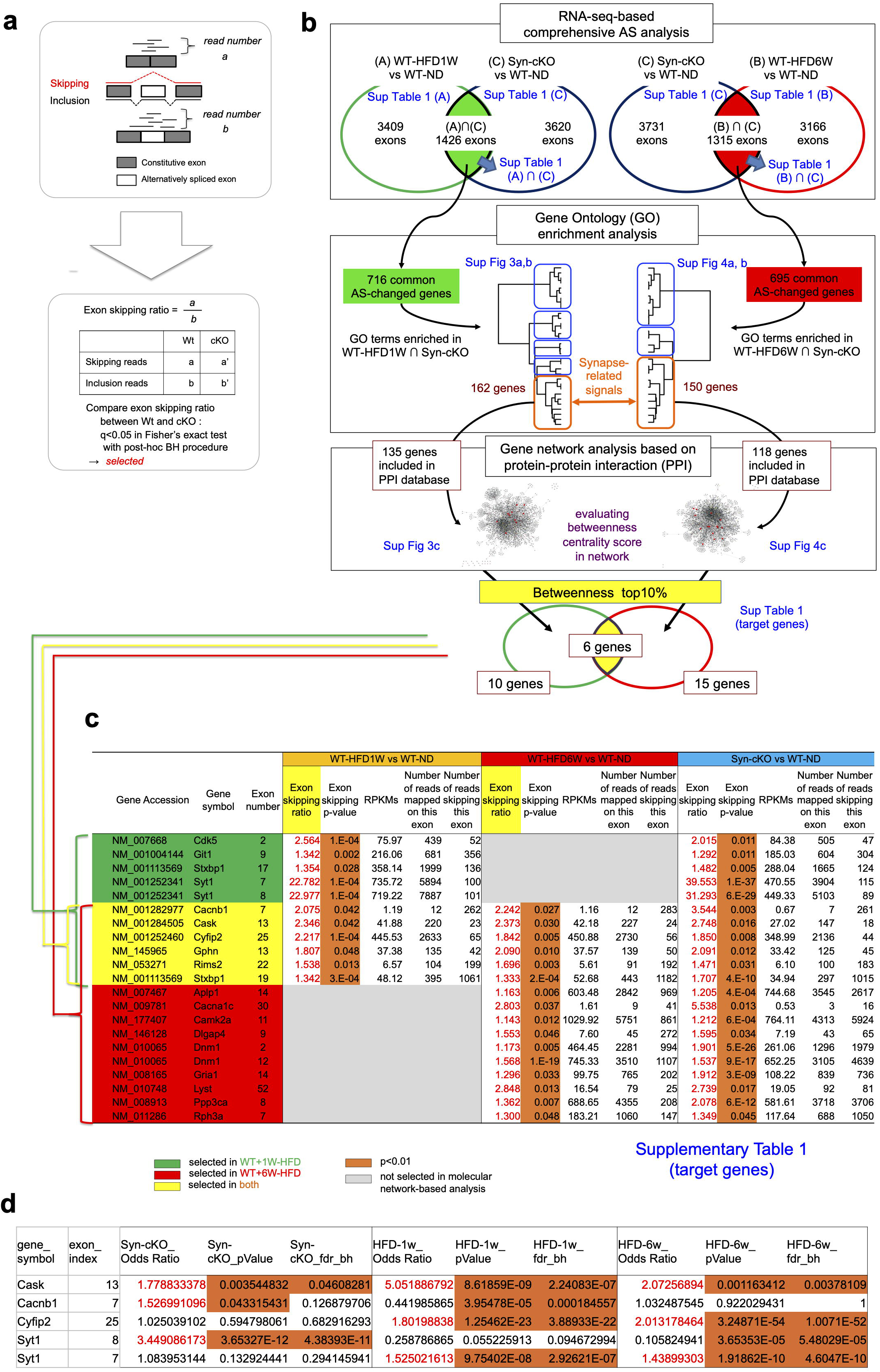
Identification of key genes by comprehensive AS analysis. a) The method for calculating the exon-skipping ratio, which was used for RNA-seq based comprehensive AS analysis.saralab) The flow of experiments and results to uncover key genes whose abnormal AS affects neuronal synapses. Detailed data in each step are shown in the corresponding Supplementary Tables or Figures (blue).saralac) Essential data of 19 key genes finally selected are shown. Supplementary Table show the details of RNA-seq analysis data.

Regarding the short-term key target gene in HFD-1W, synaptotagmin 1 (Syt1) was considered as a candidate. Though Syt1 was not selected as a final candidate by our logical flow (algorithm) to select key genes (Figure 4b), the extent of AS change in *Syt1* exon 7 or 8 was extraordinarily large (>22-fold) in HFD-1W mice (Figure 4c); the change was reproduced in independent analyses (Figure 4d); and Syt1 was closely connected to Cask and other target genes in PPI (Supplementary Figure 6b). Syt1 is a calcium sensor essential for calcium-triggered exocytosis of synaptic vesicles^71–74^. Skipping of exon 7 and/or 8 results in a critical change of Syt1 protein structure lacking C2A domain (Supplementary Figure 9) that is essential for Ca^2+^-dependent membrane fusion of synaptic vesicles^75,76^. Exon skipping of Syt1 was no longer apparent at 6W, suggesting that PQBP1-mediated AS specifically of Syt1 was recovered somehow, irrespective of the decrease of PQBP1 protein.

To strengthen our findings, we reconfirmed the abnormal AS of Cask, Cyfip2, Cacnb1 and Syt1 in HFD and their recoveries in further experiments by qRT-PCR analysis (Supplementary Figure 10).

### Functional consequence of RNA splicing impairment in target genes

Cacnb1 is a regulatory subunit of L-type calcium channel, which is widely expressed in brain and non-brain tissues and has a high voltage threshold for long-lasting activation^77^. Pre-synaptic functions of Cacnb1 are implicated, including synaptic vesicle release from GABA neurons^78^. Cyfip2 is a component of the WAVE1 complex regulating actin polymerization^79,80^ involved in the formation and stability of post-synapse dendritic spines^81–83^ and in the control of pre-synapse synaptic vesicles^84^. Cask is a pre-synapse protein identified as binding to neurexin^85^ that mediates interaction between neurexin and actin^86^ and association between synaptic vesicles and calcium channels^87^. These notions suggested that functional changes in one of the three target proteins might impair pre-synapse functions such as pre-synapse vesicle release. To test this hypothesis, we performed imaging analysis of Synapto-pHluorin (SpH), a fusion protein of pH-sensitive GFP and synaptophysin^88,89^, whose fluorescence intensities are increased when synaptic vesicle are fused to pre-synapse cell membrane. In the SpH analysis, we examined whether co-expression of full-length or skipping form of Cask, Cyfip2, Cacnb1 or Syt1 influenced the SpH signal intensities in mouse primary neurons under potassium-induced depolarization (Figure 5a), and found that full-length, but not skipped isoforms of Cyfip2 Cacnb1 or Syt1 could enhance pre-synapse vesicle release, while skipped – but not full length – isoforms of Cask could suppress pre-synapse vesicle release (Figure 5b). The results were plausible given that Cacnb1 and Syt1 are direct regulators of pre-synapse vesicle release^90,91^, and as Cyfip2 may regulate presynaptic activity like Cyfip1^92,93^ in addition to its role in post-synaptic scaffold^94^, while Cask stabilizes and maintains the synapse as a submembrane scaffold component in the pre-synapse that regulates neurexins^85^ and/or Syndecan-2^95^. These results suggested that the HFD-induced AS of Cask, Cyfip2 and Cacnb1 might impair the critical functions of pre-synapse.

**Figure 5:**
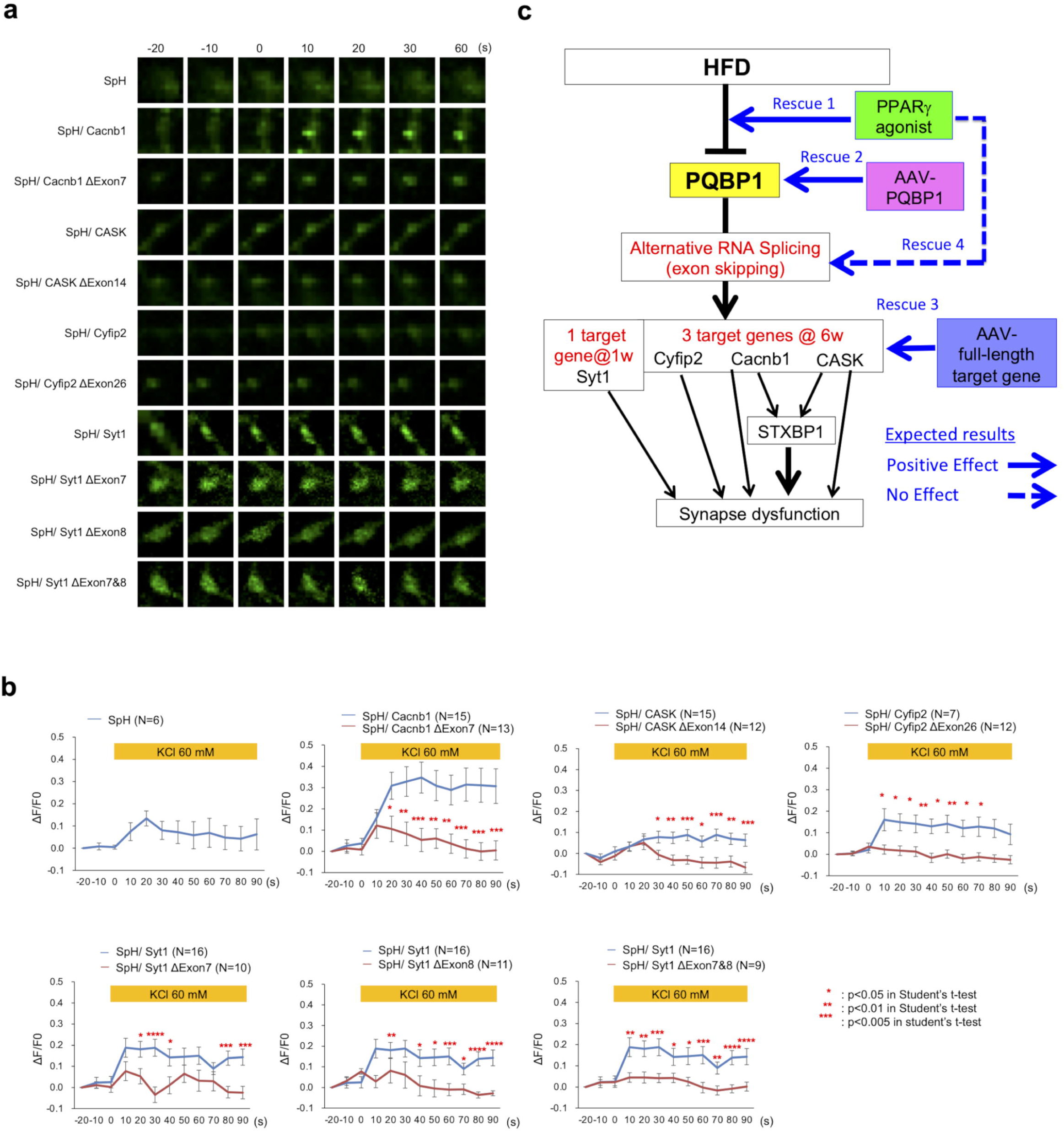
Abnormal AS of key genes impairs synapse function. a) Mouse primary cortical neurons transfected by pBI-SpH, pBI-SpH-hCacnb1, pBI-SpH-hCacnb1Δexon7, pBI-spH-hCASK, pBI-spH-hCASKΔexon14, pBI-SpH-hCyfip2, or pBI-SpH-hCyfip2Δexon26 were stimulated with 60 mM KCl and synapse depolarization was observed by time-lapse imaging of punctate-like structures close to the dendritic shaft.saralab) Intensities of SpH fluorescence at the points of −30, −20, and −10 s were averaged and used as baseline intensity (F). Next, intensities of SpH fluorescence at each time point was subtracted with the baseline intensity (ΔF). Then, the intensity was normalized with the baseline intensity and shown in the graph (ΔF/F). Statistical analyses were performed using Student’s t-test.saralac) Hypothetical scheme of the molecular pathway from HFD to synapse dysfunction via PQBP1 and AS target genes, and plan of rescue experiments to prove the hypothesis.

Collectively with the AS changes revealed by comprehensive exon-skipping analyses of HFD-induced/PQBP1-mediated AS changes (Figure 4) and the functional changes in such abnormal AS of three target genes (Figure 5), we generated our hypothesis for the molecular mechanism of synapse dysfunction under HFD (Figure 5c) and decided to test this hypothesis via in vivo experiments with PPARγ agonist, AAV-PQBP1 and AAV-full-length target genes (Figure 5c).

### PPAR**γ** agonist recovers cognitive impairment by HFD

We began by performing intra-peritoneal administration of pioglitazone, a PPARγ agonist, to HFD-6W mice or ND control mice for 3 week or 6 week duration (Figure 6a), and we investigated cognitive function in behavioral tests (Figure 6b, c, d) and evaluated synapse morphology by western blot (Figure 6e) and immunohistochemistry of cerebral cortex tissues (Figure 6f, g). As expected, administration of pioglitazone for both 3 and 6 weeks recovered long-term spatial memory on the Morris water maze test in HFD-6W mice in comparison to HFD-6W mice receiving intra-peritoneal administration of phosphate buffered saline (PBS) for 6 weeks (Figure 6b). Administration of pioglitazone for either 3 or 6 weeks also recovered short-term spatial memory in the Y-maze test (Figure 6c) and in aversive emotional memory in the fear-conditioning test (Figure 6d) in the same comparisons.

**Figure 6.**
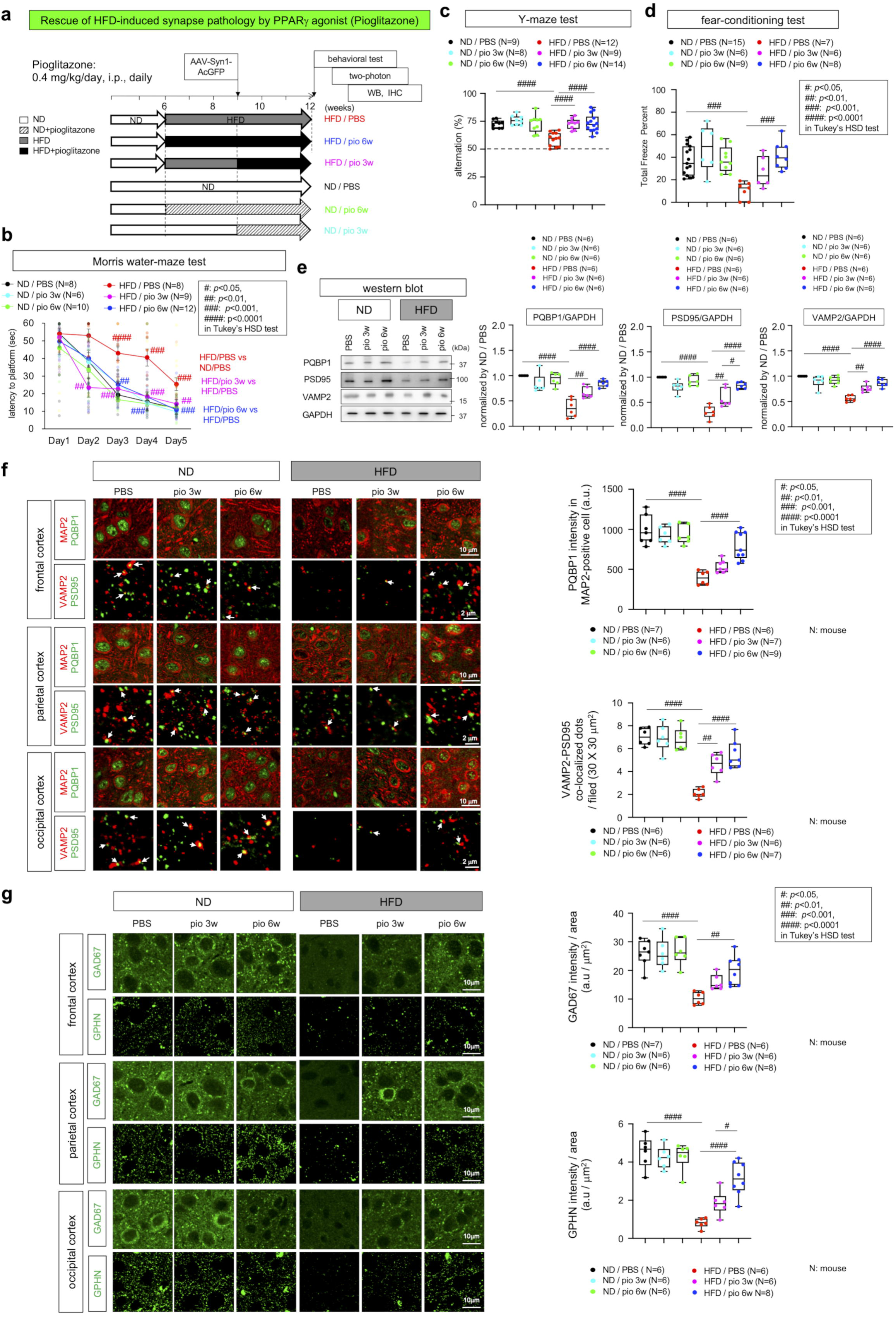
Rescue of HFD-induced synapse pathology by PPARγ agonist. a) Protocol of the rescue experiment for HFD-induced synapse pathology by pioglitazone. 6 mouse groups were prepared.saralab) Results of Morris water maze-test in the 6 mouse groups.saralac) Results of Y maze-test in the 6 mouse groups.saralad) Results of fear conditioning test in the 6 mouse groups.saralae) Western blot analysis of PQBP1, PSD95 (post-synapse marker) and VAMP2 (pre-synapse marker) in total cerebral cortex tissues of the 6 mouse groups. Right graphs show quantitative analyses of the results.saralaf) Immunohistochemistry of PQBP1 in MAP2-positive neurons and of VAMP2-PSD95 co-staining for mature synapses in 6 mouse groups. Right graphs show quantitative analyses of signal intensities of neuronal PQBP1 and of numbers of mature synapses. The mean value of three cortex areas was used as a representative value for a mouse.saralag) Immunohistochemistry of GAD67 (pre-synapse marker of inhibitory synapses) and GPHN (post-synapse marker of inhibitory synapses). Right graphs show quantitative analyses of signal intensities.

In agreement with the behavioral testing results, western blot of cerebral cortex tissues revealed recovery of PSD95 and VAMP2 expression levels in HFD-6W mice receiving pioglitzaone (Figure 6e). Moreover, immunohistochemistry of cerebral cortex revealed a decrease of VAMP2-PSD95 co-localization reflecting mature synapses in HFD-6W mice (Figure 6f), which was rescued by administration of pioglitazone for 3 weeks or 6 weeks (Figure 6f). In addition to excitatory synapses, immunohistochemistry of inhibitory synapse markers, such as GAD67 or GPNH, revealed a decrease of inhibitory synapses in the cerebral cortex of HFD-6W mice (Figure 6g), which was rescued by administration of pioglitazone for 3 or 6 weeks (Figure 6g).

### PQBP1 recovers HFD-induced cognitive impairment

We next performed intrathecal administration of AAV-PQBP1 to HFD-6W mice for three weeks before the end point (Figure 7a), and investigated their cognitive functions on behavioral tests (Figure 7b, c, d) and again evaluated synapse morphology by western blot (Figure 7e) and immunohistochemistry analysis of cerebral cortex (Figure 7f, g). As expected, AAV-PQBP1 gene therapy recovered long-term spatial memory on the Morris water maze test (Figure 7b), short-term spatial memory in the Y-maze test (Figure 7c), and aversive emotional memory in the fear-conditioning test (Figure 7d) in HFD-6W mice in comparison to HFD-6W mice receiving AAV vector only (Figure 7b, c, d). Western blot analysis of PSD95 and VAMP2 suggested recovery of excitatory synapses in the cerebral cortex of HFD-6W mice receiving AAV-PQBP1 gene therapy (Figure 7e), and immunohistochemistry of cerebral cortex revealed rescue of mature synapse decreases in HFD-6W mice receiving AAV-PQBP1 (Figure 7f). Similarly, immunohistochemistry of GAD67 and GPNH revealed rescue of inhibitory synapse numbers in HFD-6W mice receiving AAV-PQBP1 (Figure 7g).

**Figure 7:**
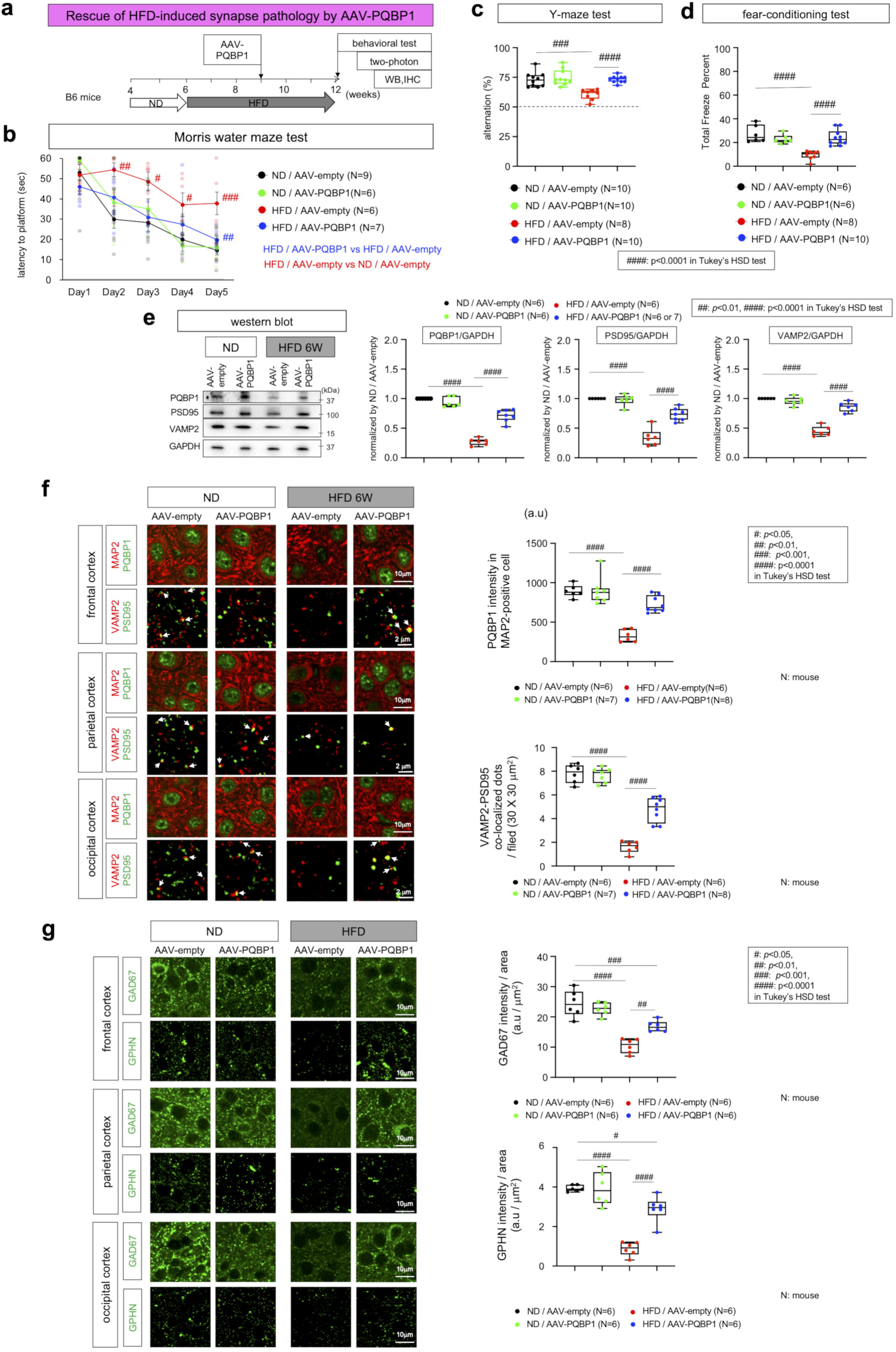
Rescue of HFD-induced synapse pathology by AAV-PQBP1. a) Protocol of the rescue experiment for HFD-induced synapse pathology by AAAV-PQBP1. 4 mouse groups were prepared.saralab) Results of Morris water maze-test in the 4 mouse groups.saralac) Results of Y maze-test in the 4 mouse groups.saralad) Results of fear conditioning test in the 4 mouse groups.saralae) Western blot analysis of PQBP1, PSD95 (post-synapse marker) and VAMP2 (pre-synapse marker) in total cerebral cortex tissues of the 4 mouse groups. Right graphs show quantitative analyses of the results.saralaf) Immunohistochemistry of PQBP1 in MAP2-positive neurons and of VAMP2-PSD95 co-staining for mature synapses in 4 mouse groups. Right graphs show quantitative analyses of signal intensities of neuronal PQBP1 and of numbers of mature synapses. The mean value of three cortex areas was used as a representative value for a mouse.saralag) Immunohistochemistry of GAD67 (pre-synapse marker of inhibitory synapses) and GPHN (post-synapse marker of inhibitory synapses). Right graphs show quantitative analyses of signal intensities.

### Restoration of normal expression of HFD-induced AS target genes can rescue synapse and cognitive impairments

We also performed intrathecal administration of AAV-Cask, -Cyfip2 or -Cacnb1 to HFD-6W mice three weeks before the end point (Figure 8a), and investigated their cognitive functions in behavioral tests (Figure 8b, c, d) and their synapse morphology by western blot (Figure 8e) and immunohistochemistry analysis of cerebral cortex tissues (Figure 8f, g).

**Figure 8:**
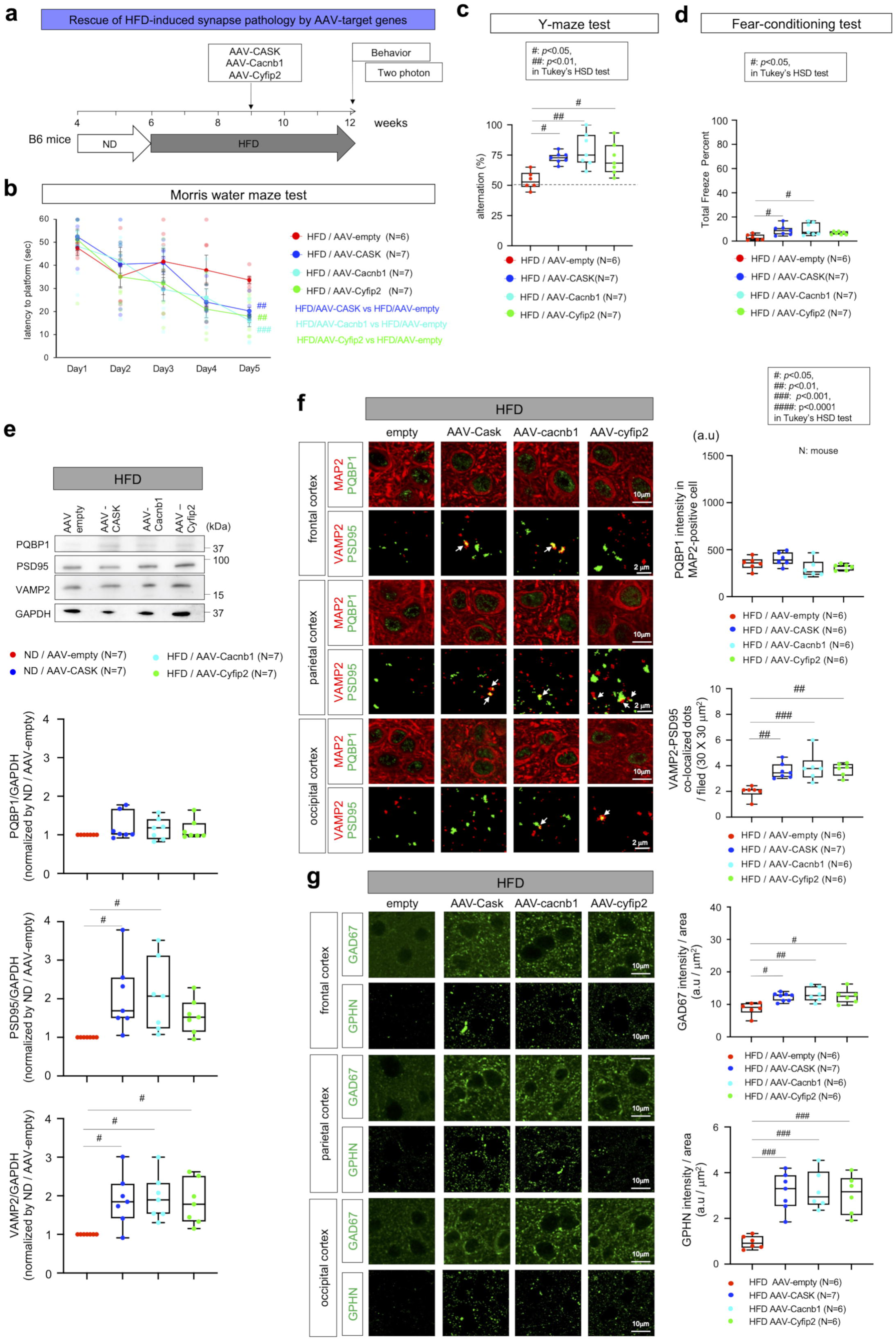
Rescue of HFD-induced synapse pathology by target genes. a) Protocol of the rescue experiment for HFD-induced synapse pathology by AAV-Cask, AAV-Cyfip2, AAV-Cacnb1 or AAV-empty. 4 mouse groups were prepared.saralab) Results of Morris water maze-test in the 4 mouse groups.saralac) Results of Y maze-test in the 4 mouse groups.saralad) Results of fear conditioning test in the 4 mouse groups.saralae) Western blot analysis of PQBP1, PSD95 (post-synapse marker) and VAMP2 (pre-synapse marker) in total cerebral cortex tissues of the 4 mouse groups. Right graphs show quantitative analyses of the results.saralaf) Immunohistochemistry of PQBP1 in MAP2-positive neurons and of VAMP2-PSD95 co-staining for mature synapses in 4 mouse groups. Right graphs show quantitative analyses of signal intensities of neuronal PQBP1 and of numbers of mature synapses. The mean value of three cortex areas was used as a representative value for a mouse.saralag) Immunohistochemistry of GAD67 (pre-synapse marker of inhibitory synapses) and GPHN (post-synapse marker of inhibitory synapses). Right graphs show quantitative analyses of signal intensities.

As expected, gene therapy of the full length version of these three target genes by AAV-Cask, AAV-Cyfip2 or AAV-Cacnb1 restored long-term spatial memory in the Morris water maze (Figure 8b), short-term spatial memory in Y-maze test (Figure 8c), and aversive emotional memory in a fear-conditioning test (Figure 8d) in HFD-6W mice in comparison to HFD-6W mice receiving AAV empty vector (Figure 8b, c, d). Western blot analysis of PSD95 and VAMP2 indicated rescue of excitatory synapses in cerebral cortex by gene therapy in HFD-6W mice (Figure 8e). Immunohistochemistry of cerebral cortex also revealed recovery of decreased mature synapses in HFD-6W mice by AAV-Cask, -Cyfip2 or -Cacnb1 (Figure 8f). Similarly, immunohistochemistry of GAD67 and GPNH indicated rescue of inhibitory synapses by all three genes (Figure 8g). AAV-Cask, -Cyfip2 or -Cacnb1 did not change the level of PQBP1 in the nuclei of cortical neurons (Figure 8e). However, improvements in behavioral tests and immunohistochemical analyses in mice receiving target gene supplementation (Figure 8) were less than that achieved by AAV-PQBP1 delivery (Figure 7). Furthermore, the therapeutic effect of AAV-Cacnb1 was much greater than that of AAV-Cyfip2 or AAV-Cask (Figure 8b, c, d, e, f, g).

### PPAR**γ** agonist does not recover synapse and cognitive impairment of PQBP1-cKO mice

Further to confirm the role of PQBP1 in HFD-induced impairment of synapse and cognitive function, we examined the effect of pioglitazone on synapse and cognitive function in Syn-cKO mice (Supplementary Figure 11). Pioglitazone should recover synapse and cognitive impairment of Syn-cKO mice, if molecules other than PQBP1 mediate the therapeutic effect of pioglitazone in previous experiments (Figure 6). Four mouse groups (Syn-cKO mice with or without pioglitazone administration, normal sibling mice with or without pioglitazone administration) were prepared for this purpose, and pioglitazone was administered from 6 to 12 weeks of age (Supplementary Figure 11a), just like the treatment for HFD-6W mice (Figure 6).

As expected, pioglitazone treatment did not recover long-term spatial memory in the Morris water maze (Supplementary Figure 11b), short-term spatial memory in the Y-maze test (Supplementary Figure 11c), or aversive emotional memory in a fear-conditioning test of Syn-cKO mice at 12 weeks of age (Supplementary Figure 11d). Western blot analysis of PSD95 and VAMP2 (Supplementary Figure 11e) as well as immunohistochemistry of cerebral cortex revealed that pioglitazone treatment of Syn-cKO mice did not improve excitatory and inhibitory synapse numbers (Supplementary Figure 11f, g). In all rescue experiments with pioglitazone, AAV-PQBP1 and AAV-target genes, we alos performed two-photon microscopy analysis and confirmed therapeutic effects on in vivo morphology of excitatory synapse in living mice (Supplementary Figure 12a, b). Taken together, these results indicate that PQBP1 is acting downstream of PPARγ and upstream of target molecules, including Cask, Cyfip2 and Cacnb1 (Figure 5c).

### PPAR**γ** agonist and Syt1 rescue HFD-induced the short-term pathology

Similar to the rescue experiments of HFD-6W, we finally examined whether intra-peritoneal administration of pioglitazone and intrathecal administration of AAV-Syt1 or AAV-PQBP1 could rescue synapse pathology in HFD-1W mice (Supplementary Figure 13, 14). As for rescue with pioglitazone, we employed pre-HFD treament or simultaneous administration with HFD (Supplementary Figure 13a), and we observed recovery of HFD-1W mice in short-tem spatial memory (Y-maze test) and emotional memory (fear-conditioning test), but remarkably not on the long-term spatial memory task (Morris water maze test) (Supplementary Figure 13b, c, d). Pre- and post-synapse marker proteins were restored by pioglitazone based upon western blot and immunohistochemistry analysis (Supplementary Figure 13e, f, g). AAV-PQBP1 administrated 2 weeks prior to the HFD-1W regimen (Supplementary Figure 14a) recovered three types of memories (Supplementary Figure 14b, c, d) and pre- and post-synapse marker proteins (Supplementary Figure 14e, f, g). These results indicate that mechanisms of synapse pathology were basically similar in the HFD-1W mice and in the HFD-6W mice, and the recovery of synapse dysfunction in HFD-1W mice involved PPARγ agonist and Syt1, as expected (Figure 5c).

## Discussion

Various mechanisms have been suggested to explain cognitive impairment and synapse dysfunction induced by HFD or high calorie diet^1–12^. However, some of the previous hypotheses were based on non-cell-autonomous pathways involving microglia^4^ or neuroinflammation^8–10^, and did not consider cell-autonomous mechanisms of neurons. Here, we discovered that PQBP1, a regulator of synapse genes in AS as well as a causative gene for human intellectual disability, is suppressed by HFD in PPARγ-mediated transcription. PPARγ is deactivated by ERK-MEK phosphorylation downstream of the insulin signaling pathway, which is activated by HFD^69^. HFD in our experiments increases blood insulin, blood glucose and body weight (Supplementary Figure 1), and the ERK-MEK-PPAR pathway transcriptionally controls expression of the *PQBP1* gene (Figure 2, Supplementary Figure 2, 3). The reduction of PQBP1 in neurons, as we expected, influences AS of synapse-related genes (Figure 4) and finally impairs excitatory and inhibitory synapses (Figure 3, 6). These abnormal changes of synapses are detected not only by immunohistochemistry but also by two-photon microscopic imaging in living animals (Figure 3, Supplementary Figure 12, 13). RNAseq analysis reveals altered expression of three key target genes (Cask, Cyfip2 and Cacnb1) in PQBP1-mediated AS (Figure 4, Supplementary Figure 4, 5) resulting in synapse dysfunction (Figure 5). In addition, Cask and Cacnb1 interact with another key gene STXBP1 (Figure 4, Supplementary Figure 6). Genetic and pharmacological approaches employed in *in vivo* rescue experiments of this study revealed that the pathway from PQBP1 to these target synapse genes controls synapse dysfunction by HFD (Figure 6, 7, 8, Supplementary Figure 11, 12, 13, 14). Though PQBP1 is involved in the pro-inflammatory cGAS-STING pathway of innate immune cells^32–34^, we did not observe expression changes of PQBP1 in microglia of HFD-mice.

From the aspect of E-I imbalance, our data revealed that HFD impairs both excitatory and inhibitory synapses (Figure 6, 7, 8, Supplementary Figure 11, 12, 13, 14). AAV-PQBP1, AAV-Cyfip2, AAV-Cacnb1, AAV-CASK or AAV-Syt1 for supplementation of the ordinary AS isoform rescues synapse and/or cognitive dysfunctions in HFD mice, supporting the pathological axes of PQBP1-Cacnb1/CASK-STXBP1, PQBP1-Cyfip2 in HFD or PQBP1-Syt1. However, the extent of synapse recovery varied among the four target gene interventions. Recovery in the excitatory synapse number by a single target molecule among three molecules was less than that of PQBP1 (Figure 7, 8), suggesting that the effect of PQBP1 on excitatory synapses are due to multiple target molecules, among which CASK, Cacnb1 and Cyfip2 display relatively large effects. The diminished recovery effect by a PQBP1 target gene also applied to Syt1 in the short-term HFD effect.

Transcriptional suppression of PQBP1 by HFD discovered in this study is consistent with the observation that HFD is a risk factor for AD^1,2^, since a previous study revealed that PQBP1 is reduced in neuronal nuclei in AD pathology^30^. At the same time, the relationship between PQBP1 in neurons and fat volume in the body has been controversial. In the brain, HFD reduces PQBP1, while PQBP1 might change differently in the periphery in response to HFD as PQBP1-deletion promotes lean body mass in patients^53^ and HFD leads to an increase of fat. In adipocytes, fatty acids not removed from the blood are ligands for PPARα and their metabolites, such as prostaglandins or leukotrienes, function as ligands for PPARγ, indicating that the short-term increase of fatty acids in HFD activates PPAR-mediated transcription. However, long-term increase of fatty acids in HFD triggers insulin release and activates ERK-MEK to deactivate PPAR-mediated transcription^96^. Moreover, this longer and tonic insulinemia would lead to insulin resistance associated with reduced PPAR activity, as supported by a number of reports^97,98^. In this regard, longer observation of changes of PQBP1 in HFD-fed animals would be interesting in order to understand the role of inter-organ communication between brain (neurons) and fat tissue (adipocytes), though animal models suitable for this type of research would ideally be primates.

Pioglitazone can cross the blood-brain barrier^99^, and functions as an agonist to the heterodimer complex of PPARγ and retinoid X receptor (RXR)^100^. In addition to its function in transcription regulation by PPARγ, pioglitazone directly or indirectly activates peroxisome proliferator activated receptor gamma coactivator-1 alpha (PGC-1α), which regulates mitochondrial function^101^. In this regard, the therapeutic effect of pioglitazone might be derived not only from recovery of PQBP1 expression, but also improved mitochondrial function. However, the failure of recovery of Syn-cKO mice treated with pioglitazone (Figure 9) indicates that contribution of such a PQBP1-independent mechanism is minor in the therapeutic effect of pioglitazone.

Pioglitazone has been tested in clinical trials as a treatment for cognitive impairment in AD at-risk individuals based upon a biomarker risk assignment algorithm (BRAA)^102^ and in patient with DM type 2 and ischemic stroke who are at risk for developing dementia^103^, and the results were respectively negative and positive. As the BRAA criteria in the former clinical trial employed ApoE and TOMM40 genotypes, the cohort might not exactly match individuals with HFD or high blood glucose. Therefore, further assessment of the potential beneficial effects of this PPARγ agonist for HFD-associated dementia or HFD-associated AD is necessary.

Considering that PQBP1 dysfunction could occur in neurons with AD pathology, where PQBP1 is reduced due to depletion of the scaffold protein SRRM2 for PQBP1 in the nucleus^30^, pioglitazone may not be effective in HFD-associated AD patients who already have progressed to advanced PQBP1 dysfunction in the brain. Thus, early administration of pioglitazone to achieve maximum therapeutic benefit might be necessary. In such cases of advanced HDF-induced AD, AAV-PQBP1 gene therapy to boost PQBP1 expresison levels could be a viable option for future therapy development. Even though previous data indicate that AAV-PQBP1 driven by the CMV promoter increases PQBP1 selectively in neurons but not in microglia^30,34^, it will also be necessary to select an enhancer/promoter that will restrict PQBP1 expression to neurons in the central nervous system, because of the potential role of PQBP1 in oncogenesis and cancer progression^31^.

## Methods

### Plasmid construction

Full-length human cDNA of Cask, Cacnb1 or Cyfip2, which were amplified from HeLa cell mRNA by PCR with the following primers, were subcloned between Mlu I and Not I sites of pCI-neo plasmid (Promega, E1841, Madison, WI, USA) in which 3xMyc-tag was added at C-terminal in reading frame. From these plasmids expressing full-length mRNA, pCI-neo plasmids expressing exon-skipped Cask(Δexon 14), Cacnb1(Δexon 7) and Cyfip2(Δexon 25) were generated by the following PCR primers.

An expression vector for full length cDNA of mouse STXBP1 with His-tag (pCMV3-His-mSTXBP1) was purchased from Sino Biological (MG58001-NH, Beijing, China).

For imaging analysis of pre-synapse vesicle release, full-length Cask, Cacnb1, Cyfip2 or exon-skipped Cask(Δexon 14), Cacnb1(Δexon 7), Cyfip2(Δexon 25) cDNA was subcloned between Mlu I and Not I sites of a bidirectional expression vector pBI-SpH in which rat synapto-pHluorin fused to rat VAMP2 had been already subcloned. The pBI-SpH was generated by subcloning rat VAMP2 cDNA between EcoR I and Xba I and then subcloning rat synapto-pHluorin cDNA amplified from CMV::SypHy A4 (Addgene, #24478, Watertown, MA. USA) into pBI-CMV1 (Clontech, #631630, Mountain View, CA, US) at Age I site included by the PCR primer for amplification of rat VAMP2 cDNA.

Full-length cDNA of mouse Syt1, which was purchased from Dnaform (Yokohama, Japan) as RIKEN FANTOM clone 8030489K02 (Wako, Japan), was subcloned into pBI-CMV1 (Clontech, Cat# 631630, Mountain View, CA, US) containing synapto-pHluorin cDNA by NEBuilder HiFi DNA Assembly Master Mix Kit (BioLabs, Cat# E2621S/L/X, MA, USA), and expression vectors for Δexon 7, Δexon 8 and Δexon 7&8 were generated by PCR using the following primers.

The primer sequences were:

Cask Forward: CCACGCGTATGGCCGACGACGACGT

Cask Reverse: GGGCGGCCGCCTCGAGCTAATAGACCCAGGAGACA

Cask(Δexon 14) Forward: GCCAAAGAGGCCTACTTCAGACTCAC

Cask(Δexon 14) Reverse: AAGTAAGGCCTCTTTGGCTCTCTTGTAC

Cacnb1 Forward: TGACGCGTATGGTCCAGAAGACCAGCATGTCC

Cacnb1 Reverse: ATGCGGCCGCGAATTCTCAGCGAATGTAGACGCC

Cacnb1(Δexon 7) Forward: CTGCCAGTGACAGAGCATGTGCCCCC

Cacnb1(Δexon 7) Reverse: CATGCTCTGTCACTGGCAGGGGGTGTG

Cyfip2 Forward: CCACGCGTGTCGACATGACCACGCGACGTCACCCTGG

Cyfip2 Reverse: GGGCGGCCGCCTCGAGTTAGCAAGTGGTGGCAAGGAC

Cyfip2 (Δexon 25) Forward: GTGAAGAGCTTGGGATCCTGGAGTTCTTCC

Cyfip2 (Δexon 25) Reverse: GAACTCCAGGATCCCAAGCTCTTCAATCTTTA

Syt1 Forward: ATCCGCTAGGGATCCATGGTGAGTGCCAGT

Syt1 Reverse: TGCGGCCGCGCTAGCTTACTTCTTGACAGC

Syt1(Δexon 7) Forward: ATGCCTGCTGGTGGGAATC

Syt1(Δexon 7) Reverse: ATGAAGGATCAGCTGCTGGTGGGAATCATC

Syt1(Δexon 8) Forward: CTAATTCCGAGTATGGCACCTGGTTATTCTGGAA

Syt1(Δexon 8) Reverse: TTCCAGAATAACCAGGTGCCATTACTCGGAATTAG

Syt1(Δexon 7&8) Forward: CTAATTCCGAGTATGGCACCTGATCCTTCAT

Syt1(Δexon 7&8) Reverse: ATGAAGGATCAGGTGCCATACTCGGAATTAG

VAMP2 Forward: TCGAATTCTGATGTCGGCTACCGCTGC

VAMP2 Reverse: GGTCTAGATTAACCGGTCTGCTGAAGTAAACGATGA

All the plasmids after constructions were verified by DNA sequencing.

### Generation of Pqbp1-cKO mice

Pqbp1-floxed heterozygous or homozygous female mice^27^ were crossed with Synapsin1-Cre transgenic heterozygous male mice (The Jackson Laboratory, Bar Harbor, ME, USA) to generate Pqbp1 conditional knockout mice, as we reported previously^27^.

The genotyping of pups was performed by PCR with primer as listed below; Floxed allele: wild-type (499 bp), floxed (539 bp)

5’-AATCTTGGAGTTAGTAATGGTGCTT,

3’-AATCTCATGTAATTGACGAGACAGAG

Synapsin1 promoter-Cre: transgene (300 bp)

5’-CTCAGCGCTGCCTCAGTCT

3’-GCATCGACCGGTAATGCA

Synapsin1 promoter-Cre: internal positive control (206 bp)

5‘-CTAGGCCACAGAATTGAAAGATCT

3’-GTAGGTGGAAATTCTAGCATCATCC

Pqbp1-cKO mice (Syn-cKO) were judged from the PCR products’ band size.

### Animal and Diet

Male C57BL/6J mice (Sankyo Labo Service Corporation, INC, Japan) were caged in a temperature-controlled room (25°C) and exposed to daily 12 h light-12 h dark cycle with free access to water and food. The mice were fed with normal diet (CLEA Rodent Diet CE-2, CELA, Japan) or HFD (High Fat Diet 32, CELA, Japan) according to the protocol indicated in Figures. Blood glucose was measured by using Glucose Pilot (NGP-01B, Technicon international, Tokyo, Japan) weekly. Plasma insulin levels were measured by ELISA kit (Mouse Insulin ELISA, Mercodia, Uppsala, Sweden) according to the manufacturer’s protocol.

### RNA sequencing

Mouse brain tissue samples were collected and homogenized in 350□µL RNA RLT buffer (Qiagen)/0.01% 2-mercaptoethanol (Wako, Tokyo, Japan). Total RNA was purified with RNeasy mini kit (Qiagen). To eliminate genomic DNA contamination, on-column DNA digestion was conducted for each sample with DNase I (Qiagen). Prepared RNA samples were subjected to a HiSeq-based RNA-seq by TAKARA (700 million bp reads).

### Analysis of alternative splicing changes

Alternative splicing profiles of each sample were evaluated by the number of short reads that were mapped onto or skipped exons in the reference mouse genome assembly mm39. For each exon, Fisher’s exact test was employed to evaluate the statistical significance of changes by comparing “mapped” and “skipping” counts between two groups. To increase the reliability of this analysis, exons with no more than 10 “mapped” or “skipping” counts in both groups were excluded.

### GO enrichment analysis

Gene Ontology (GO) enrichment analysis was performed using DAVID (https://david.ncifcrf.gov/home.jsp). The enriched GO terms were classified by hierarchical clustering using Ward’s method. To calculate the distance between GO terms in Ward’s method, the associated gene lists of them were used.

### Network centrality analysis of the pathological network

To generate the pathological network based on Protein-Protein interaction (PPI), UniProt accession numbers were added to genes identified in RNA-seq-based alternative splicing (AS) analysis. The pathological PPI network was constructed by connecting genes using the integrated database of the Genome Network Project (GNP) (https://cell-innovation.nig.ac.jp/GNP/index_e.html), which includes BIND, BioGrid (http://www.thebiogrid.org/), HPRD, IntAct (http://www.ebi.ac.uk/intact/site/index.jsf), and MINT. A database of GNP-collected information was created on the Supercomputer System available at the Human Genome Center of the University of Tokyo.

The network was expanded to one-hop (directly linked) connections from AS-changed gene’s nodes, and the degree of significance of AS-changed genes were evaluated by calculating betweenness centrality score. Genes having top 10% of betweenness score were considered as important nodes in the pathological network.

### Rescue experiments

In the pharmacological rescue experiment, 0.4 mg/kg pioglitazone hydrochloride (Wako, Cat#168-24833, Osaka, Japan) in 100 μl PBS was intraperitoneally injected to mice every day. In the genetic rescue experiments, AAV1-PQBP1 with the Cytomegalovirus (CMV) promoter^27^ (titer: 1□×□10^8^ vector genomes/μL, 1□μL), AAV9-Cask with the Cytomegalovirus (CMV) promoter (titer: 1□×□10^8^ vector genomes/μL, 1□μL, VectorBuilder, Burlingame, CA, USA), AAV9-Cacnb1 with the Cytomegalovirus (CMV) promoter (titer: 1□×□10^8^ vector genomes/μL, 1□μL, VectorBuilder, Burlingame, CA, USA), AAV9-Cyfip2 with the Cytomegalovirus (CMV) promoter (titer: 1□×□10^8^ vector genomes/μL, 1□μL, VectorBuilder, Burlingame, CA, USA), AAV9-Syt1 with the Cytomegalovirus (CMV) promoter (titer: 1□×□10^8^ vector genomes/μL, 1□μL, VectorBuilder, Burlingame, CA, USA), or AAV1-empty with the Cytomegalovirus (CMV) promoter (titer: 1□×□10^8^ vector genomes/μL, 1□μL, VectorBuilder, Burlingame, CA, USA) was injected to the subarachnoid space on the surface of the retrosplenial cortex at (−1.0 mm from bregma; lateral 0.5 mm) under anesthesia with 1% isoflurane.

### Mouse behavioral analysis

Spatial memory during exploratory behavior was assessed by the Y-maze test, using Y-shape maze with three identical arms interconnected at an angle of 120° (YM-3002, O’HARA & Co., Ltd, Tokyo, Japan). After mice being habituated to the procedure room at least 30□min prior to testing, they were put at the end of one arm and allowed to explore freely through the maze during an 8□min session. The percentage of spontaneous alterations (indicated as an alteration score) was calculated based on the sequence of arm entries, dividing the number of entries into a new arm different from the previous one by the total number of transfers from one arm to another.

In the Morris water maze test, mice were trained to escape from water by swimming onto a hidden platform. Water temperature was maintained at 21□±□1□°C. Mice performed four training trials per day (60□s) for 4 days, and the latency to reach the hidden platform was recorded.

In the fear-conditioning test, the freezing response was measured at 24□h after the conditioning trial (65□dB white noise, 30□s□+□foot shock, 0.1□mA, 2□s) in the same chamber in the absence of a foot shock.

### Cell culture and transfection

HeLa and HEK 293T cells were maintained at 37°C, 5% CO_2_ in Dulbecco’s Modified Eagle’s Medium–high glucose (D5796 medium, Sigma Aldrich, St. Louis, MO, USA) with 10% Fetal Bovine Serum (#10270106, Gibco, MA, USA), Cells were transfected with plasmids using Lipofectamine 2000 (Thermo Fisher Scientific, Waltham, MA, USA), according to the manufacturer’s protocol.

Mouse primary cerebral neurons were prepared from embryonic Day 15 C57BL/6J mouse embryos. Cerebral cortexes were dissected, incubated with 0.05% trypsin in 4 mL of phosphate buffered saline (PBS) (Thermo Fisher Scientific, Waltham, MA, USA) at 37□°C for 15 min, and dissociated by pipetting. The cells were passed through a 70 μm cell strainer (Thermo Fisher Scientific, Waltham, MA, USA), collected by centrifugation, and cultured in neurobasal medium (Thermo Fisher Scientific, Waltham, MA, USA) containing 2% B27, 0.5 mM L-glutamine, and 1% Penicillin/Streptomycin in the presence of 0.5 μM AraC. Immunohistochemistry of primary neurons was performed at 14 days after cell preparation.

Mouse primary microglia were prepared from cerebral cortexes of C57BL/6J mice at P1–P3. Cortex tissues were minced, and digested with 0.05% Trypsin-EDTA (#25200056, Thermo Fisher Scientific Inc., MA, USA) at 37□°C for 10□min. Trypsinization was stopped by adding 10% fetal bovine serum, and 10□mg/ml DNase was added to digest genomic DNA. The cells were collected by centrifugation at 1,000□×l*g* for 5□min, dissociated by pipetting and passage through a 40□μm cell strainer (#22-363-548, Thermo Fisher Scientific Inc., MA, USA), and seeded into a T-75 culture flask in primary microglia culture medium (MGC57, Cosmo Bio Co., Ltd, Tokyo, Japan) supplemented with 1% penicillin/streptomycin. Medium was replaced once a week. At DIV 14-21, microglial cells were isolated from confluent mixed glial cultures by shaking (80□r.p.m. for 60□min at 37□°C) and centrifuged at 1,000□g for 10□min. The floating cells were collected and seeded on 96 well plates or 12-well plates. Microglial cells were used after another 24□h.

### Luciferase assay

Culture medium of primary neuron was replaced with neurobasal medium (Thermo Fisher Scientific, Waltham, MA, USA) containing 2% B27, 0.5 mM L-glutamine, and 1% Penicillin. After 5 days, by using lipofectamine 2000 (Thermo Fisher Scientific, Waltham, MA, USA) 1□×□10^4^ primary neurons were transfected with 2.5 ng of pPQBP1 5’UTR −1.8k or pPQBP1 5’UTR −1.6k, together with 2.5 ng of pCI-PPARg, pCI-PPARa, pCI-PPARd or pCI-empty, and 2.5 ng Renilla Luciferase. After 24H of transduction, luciferase assay was performed using the Dual-Glo Luciferase assay system (Promega, #E2940, WI, USA). To examine the effect of insulin, insulin (4501-V, Peptide institute, Osaka, Japan) was added at 4 or 8 ng/ml to the medium.

### Immunohistochemistry

Mouse brains were fixed in 4% paraformaldehyde for 12□h and embedded in paraffin. Sagittal sections (thickness, 5□μm) were de-paraffinized in xylene, re-hydrated, dipped in 0.01□M citrate buffer (pH 6.0), and microwaved at 120□°C for 15□min. Sections were blocked with 10% FBS containing PBS. Immunohistochemistry was performed using the following primary antibodies: mouse anti-PQBP1 (1:400, sc-374260, Santa Cruz Biotechnology, Dallas, TX, USA); goat anti-Iba1 (1:500, 011-27991, Wako, Osaka, Japan); rabbit anti-S100B (1:500, ab52642, Abcam, Cambridge, UK); mouse anti-MAP2 (1:500, sc-32791, Santa Cruz Biotechnology, Dallas, TX, USA); mouse anti-VAMP2 (1:200, 13647S, Cell Signaling Technology, Danvers, MA, USA); rabbit anti-PSD95 (1:200, 3409S, Cell Signaling Technology, USA). mouse GAD67 (1:200, ab26116, Abcam, Cambridge, UK). mouse GPHN (1:200, ab177154, Abcam, Cambridge, UK) Secondary antibodies were as follows: Cy3-AffiniPure Donkey Anti-Goat IgG (H + L) (1:500, 705165003 Jackson ImmunoResearch); Cy3-AffiniPure Donkey Anti-Rabbit IgG (H + L) (1:500 711165152 Jackson ImmunoResearch); donkey anti-mouse IgG Alexa488 (1:1000, A21202, Molecular Probes, Eugene, OR, USA). Nuclei were stained with DAPI (0.2 μg/ml in PBS, D523, DOJINDO Laboratories, Kumamoto, Japan).

### Western blot analysis

Hela cell and HEK 293T cell in culture dishes were washed three times with PBS and dissolved in TNE buffer (10□mM Tris-HCl (pH 7.5), 150□mM NaCl, 1□mM EDTA, 1% Nonidet P-40), containing protease inhibitor cocktail (#539134, Merck Millipore). Then mixed with sample buffer (62.5□mM Tris-HCl, pH 6.8, 2% (w/v) SDS, 2.5% (v/v) 2-mercaptoethanol, 10% (v/v) glycerol, and 0.0025% (w/v) bromophenol blue.) Samples were separated by SDS-PAGE, transferred to Immobilon-P polyvinylidene difluoride membranes (Millipore, Burlington, MA, USA) using the semi-dry method, blocked with 5% milk in TBST (10□mM Tris-HCl pH 8.0, 150□mM NaCl, and 0.05% Tween-20), and incubated overnight at 4□°C with the following primary antibodies diluted in Can Get Signal solution (Toyobo, Osaka, Japan as follows: mouse anti-PQBP1 (1:3000, sc-374260, Santa Cruz Biotechnology, Dallas, TX, USA); rabbit anti-PSD95(1:3000, 3409 Cell Signaling Technology, USA); mouse anti-VAMP2 (1:3000, 13647S, Cell Signaling Technology, Danvers, MA, USA); mouse anti-GAPDH (1:5000, MAB374, Millipore, Burlington, MA, USA); rabbit anti-Cask (1:10,000, ab80579, Abcam, Cambridge, UK); mouse anti-Cacnb1 (1:3000, ab85020, Abcam, Cambridge, UK); rabbit anti-Cyfip2 (1:3000, ab95969, Abcam, Cambridge, UK). The membranes were subsequently incubated with HRP-linked anti-rabbit IgG (1:3000, #NA934, GE Healthcare) or HRP-linked anti-mouse IgG (1:3000, #NA931, GE Healthcare) secondary antibodies for 1□h at room temperature. ECL Select Western Blotting Detection Reagent (RPN2235, Cytiva, Marlborough, MA, USA) and an ImageQuant LAS 500 luminescent image analyzer were used to detect proteins. Full-scan images are displayed in Source data file.

### Immunoprecipitation

HeLa cells were cultured with Dulbecco’s Modified Eagle’s Medium–high glucose (D5796 medium, Sigma Aldrich, St. Louis, MO, USA) with 10% Fetal Bovine Serum (#10270106, Gibco, MA, USA), transfected with the plasmids of PCI-Cacnb1-3myc, PCI-Cask-3myc or pCMV-His-STXBP/Munc18-1, and harvested 48□h later.

The cells were lysed with TNE buffer (10□mM Tris-HCl (pH 7.5), 150□mM NaCl, 1□mM EDTA, 1% Nonidet P-40) containing protease inhibitor cocktail (#539134, Merck Millipore). The lysates were rotated for 60□min at 4□°C and centrifuged at 12,000□×□g□for 1□min at 4□°C. Each supernatant was incubated with a 50% slurry of Protein-G Sepharose beads (17061801, GE Healthcare, Chicago, IL, USA) for 2□h at 4□°C, followed by centrifugation at 2000□×□g□for 2□min at 4□°C. The supernatants were incubated with 2□μl antibody for 12 h at 4□°C with rotation, followed by the addition of 40 μl Protein-G Sepharose and rotation for 2 h at 4□°C. The beads were washed three times with TNE buffer, and 30□μl sample buffer (125□mM Tris-HCl (pH 6.8), 4% (w/v) SDS, 5% (v/v) 2-mercaptoethanol, 10% (v/v) glycerol, and 0.0025% (w/v) bromophenol blue) were added to each sample. The samples were boiled at 100□°C for 5□min, followed by SDS-PAGE and transfer to Immobilon-P polyvinylidene difluoride membranes (Merck Millipore). The membranes were incubated with various primary antibodies, including mouse anti-HIS antibody (#D291-3, MBL, Aichi, Japan); mouse anti-Myc antibody (#M047-3, MBL, Aichi, Japan).

### Quantitative RT-PCR

Total RNA was isolated from entorhinal cortex tissues of ND / HFD-1W / HFD-6W mice at 12 weeks with RNeasy mini kit (74106, Qiagen, Limburg, The Netherlands). To eliminate genomic DNA contamination, on-column digestion of DNA was carried out with DNase I. Reverse transcription was performed using the SuperScript VILO cDNA Synthesis kit (11754-250, Invitrogen, Carlsbad, CA, USA). Quantitative PCR analyses were performed with the 7500 Real-Time PCR System (Applied Biosystems, Foster City, CA, USA) using the Thunderbird SYBR Green (QPS-201, TOYOBO, Osaka, Japan) and assessed by the standard curve method. The primer sequences were:

mouse PQBP1, forward primer: 5′-AGGGGCATCCTCAAACATCTG-3′ and reverse primer: 5′-CAAACACCTTGTACCAGCTCG-3′

mouse GAPDH, forward primer: 5′-TGAACGGGAAGCTCACTGG-3′ and reverse primer: 5′-TCCACCACCCTGTTGCTGTA-3′

The PCR conditions for amplification were 95□°C for 10□min for enzyme activation, 95□°C for 15□s for denaturation, and 60□°C for 1□min for extension (40 cycles). The expression levels of individual genes were normalized against the expression of GAPDH and calculated as a relative expression level.

### Two-photon microscopic analysis

Injected both Adeno-associated virus 1 (AAV1)-EGFP harboring the synapsin I promoter (titer: 1□×□10^8^ vector genomes/μL; 1 μL)^30^ and AAV2-VAMP2-mCherry harboring the CMV promoter (titer: 1×10^8^ vector genomes/μL; 1 μL)^30^ to the subarachnoid space on the surface of the retrosplenial cortex at (−3.0 mm from bregma; lateral 0.5 mm) under anesthesia with 1% isoflurane.

After 4 weeks, the skull was thinned with a high-speed micro-drill in the mouse splenial cortex. Then, the head of each mouse was immobilized by attaching the head plate to a custom machine stage mounted on the microscope table. Two-photon imaging was performed using a laser-scanning microscope system FV1000MPE2 (Olympus, Tokyo, Japan) equipped with an upright microscope (BX61WI, Olympus, Japan), a water-immersion objective lens (XLPlanN25xW; numerical aperture, 1.05), and a pulsed laser (MaiTaiHP DeepSee, Spectra Physics, Santa Clara, CA, USA). EGFP and mCherry were excited at 920 nm and scanned at 495–540 nm and 575–630 nm, respectively. High-magnification imaging (101.28 μm × 101.28 μm; 1024 × 1024 pixels; 1 μm Z step) of cortical layer I was performed with a 5□×□digital zoom through a thinned-skull window in the frontal cortex. Blinded observers performed image acquisition and analysis. Image processing was performed with Imaris Interactive Microscopy Image Analysis software (Bitplane, Zurich, Switzerland).

### Imaging analysis of pre-synapse vesicle release

Mouse primary cortical neurons were prepared from embryos at E15. Ten days later, cultured neurons were transfected by pBI-SpH, together with pBI-SpH-hCacnb1, pBI-SpH-hCacnb1Δexon7, pBI-SpH-hCASK, pBI-SpH-hCASKΔexon14, pBI-SpH-hCyfip2, or pBI-SpH-hCyfip2Δexon26 using viromer RED (lipocalyx, #TT10302, Halle, Germany) at a concentration of 100 ng plasmid / 6 × 10^4^ cells / well. Twenty-four hours after transfection, neurons were stimulated with high concentration of KCl (60 mM, sigma, #24-4290-5, St. Louis, MO, USA). Time-lapse imaging was performed by Olympus FV1200 confocal microscopy (Olympus, Tokyo, Japan) using following acquisition property (objective UPLANSAPO x40, zoom 1, 512 x 512 pixels, Argon laser, 1 channel (EGFP) filter). Images were acquired from 30 seconds before KCl stimulation to 90 seconds after KCl stimulation. Punctate-like SpH structures close to the dendritic shafts were marked, and fluorescence signals of SpH were analyzed by Image J (URL: https://imagej.nih.gov/ij/). Intensities of SpH fluorescence at the points of −20s were used as baseline intensity (F). Next, intensities of SpH fluorescence at each time point was subtracted with the baseline intensity (ΔF). Then, the intensity was normalized with the baseline intensity and shown in the graph (ΔF/F). Statistical analyses were performed using Student’s t-test.

### Reverse transcription PCR

Total RNA was isolated from cortex tissues of mice with RNeasy mini kit (74106, Qiagen, Limburg, The Netherlands). To eliminate genomic DNA contamination, on-column digestion of DNA was carried out with DNase I. Reverse transcription was performed using the SuperScript VILO cDNA Synthesis kit (11754-250, Invitrogen, Carlsbad, CA, USA). The primer sequences were:

Cask Forward: TGTTTTCCAGGATCAACATC

Cask Reverse: TGGAGAATCACCGTTTAAATA

Cacnb1 Forward: TCCAGCAAGTCAGGTGAC

Cacnb1 Reverse: CTCATAGCCCTTGAGCGA

Cyfip2 Forward: TAGCTCCTACCGGAACTTT

Cyfip2 Reverse: AGAACAGGATGGCGTTGC Syt1

Forward: GCTGCTTCTGTGTCTGTAA Syt1

Reverse: TTGCCACCTAATTCCGAGT

## Ethics

This study was performed in strict accordance with the recommendations of the Guide for the Care and Use of Laboratory Animals of the Japanese Government and National Institutes of Health. All experiments were approved by the Committees on Gene Recombination Experiments, Human Ethics, and Animal Experiments of the Tokyo Medical and Dental University (G2018-082C3, O2020-002-03, 2014-5-3 and A2021-211A).

## Data availability

All data generated or analyzed during this study are included in this article.

