## Supplementary Figure 1-14 for "PQBP1-dependent alternative RNA splicing underlies high calorie diet-induced cognitive impairment"

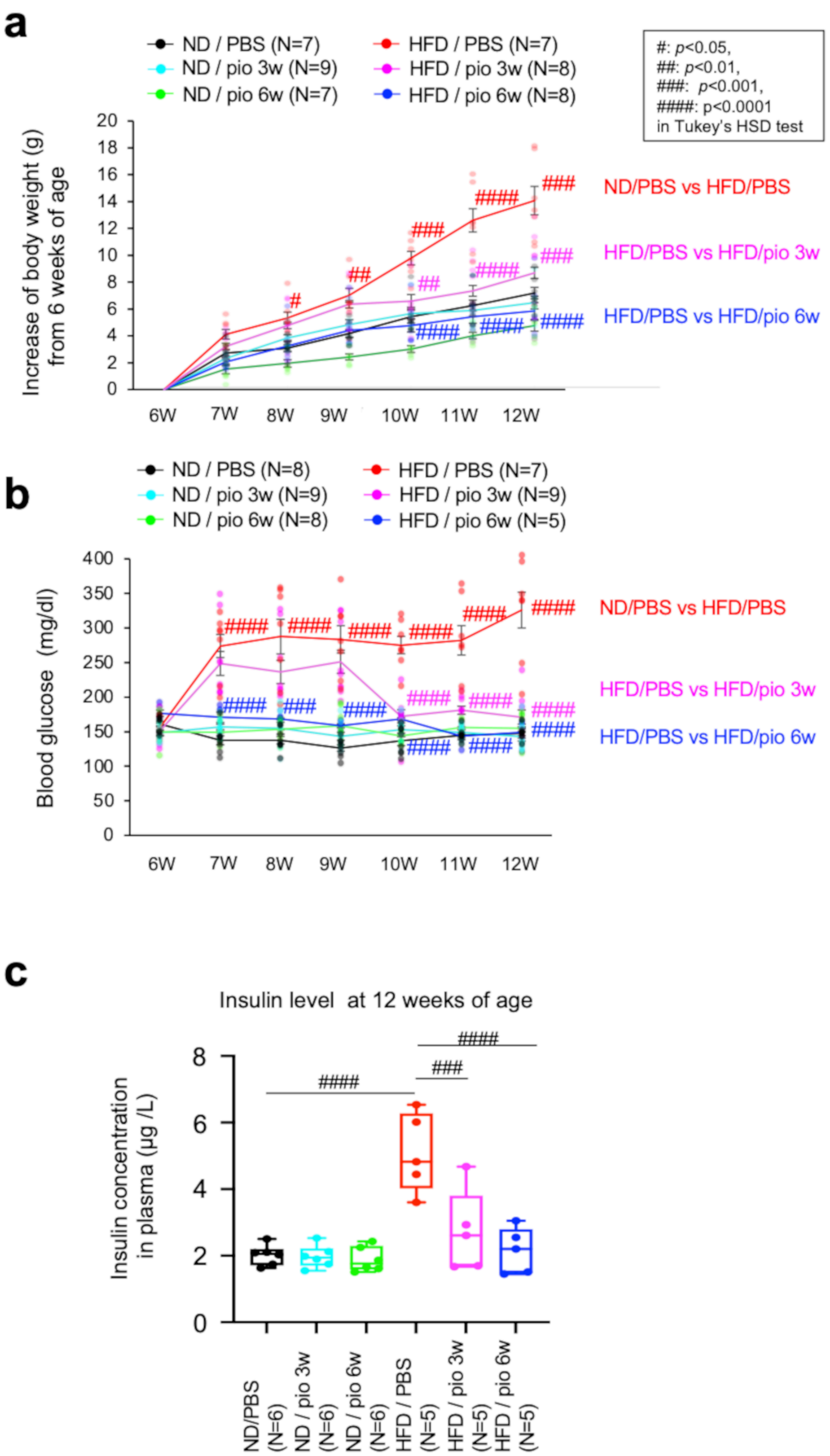


**Supplementary Figure 1**
**Clinical data of HFD mice and transcriptional regulation of PQBP1 in microglia**
a) Body weights were measured in ND and HFD mice receiving peritoneal injection of pioglitazone or PBS.
b) Blood glucose levels were measured in ND and HFD mice receiving peritoneal injection of pioglitazone or PBS.
c) Blood insulin levels were measured in ND and HFD mice receiving peritoneal injection of pioglitazone or PBS

**Supplementary Figure 2**

**a** The Human Protein Atlas (https://www.proteinatlas.org/)

| Expression level by RNA-seq (nTPM) | PPARg | PPARa | PPARd |
|---|---|---|---|
| HEK293 | 20.3 | 8.9 | 11.8 |
| HeLa | 8.1 | 4.8 | 10.9 |
| Excitatory neurons | 38.8 | 1.4 | 13.5 |
| Inhibitory neurons | 72.6 | 13.5 | 18.3 |

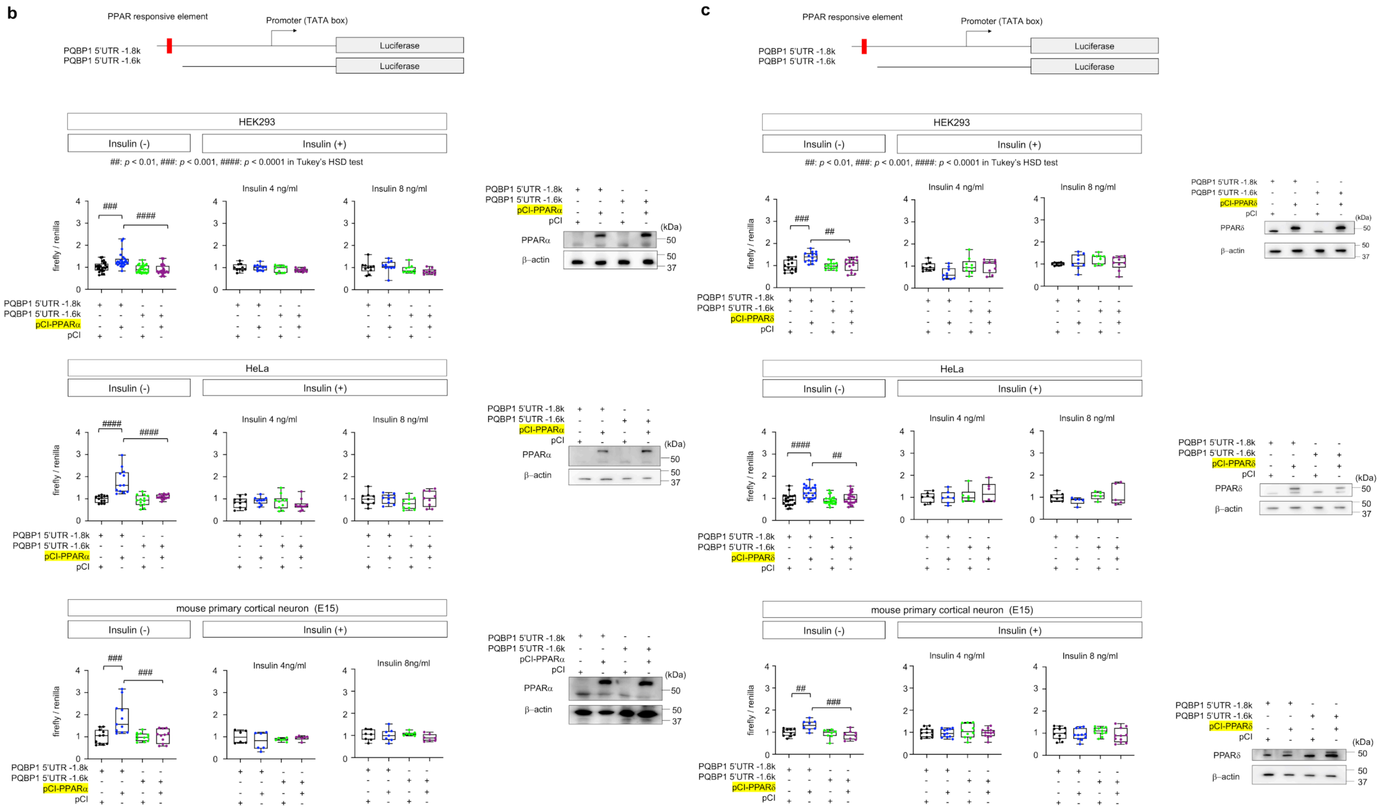


#### Supplementary Figure 2
#### PPARα and PPARδ regulate PQBP1 gene expression

a) Endogenous expression levels of PPARγ and PPARα and PPARδ according to the Human Protein Atlas database (https://www.proteinatlas.org/).

b) Reporter vectors possessing 1.8 kb or 1.6 kb of the upstream region of mouse *PQBP1* gene were used for luciferase assay.

b) Luciferase assay was performed in HEK293 cells, HeLa cells and mouse primary cortical neurons, to examine the effect of PPARα on *PQBP1* gene expression. Right panels show expression of PPAR in the absence of insulin.

c)) Luciferase assay was performed in HEK293 cells, HeLa cells and mouse primary cortical neurons, to examine the effect of PPARδ on *PQBP1* gene expression. Right panels show expression of PPAR in the absence of insulin.

### Supplementary Figure 3

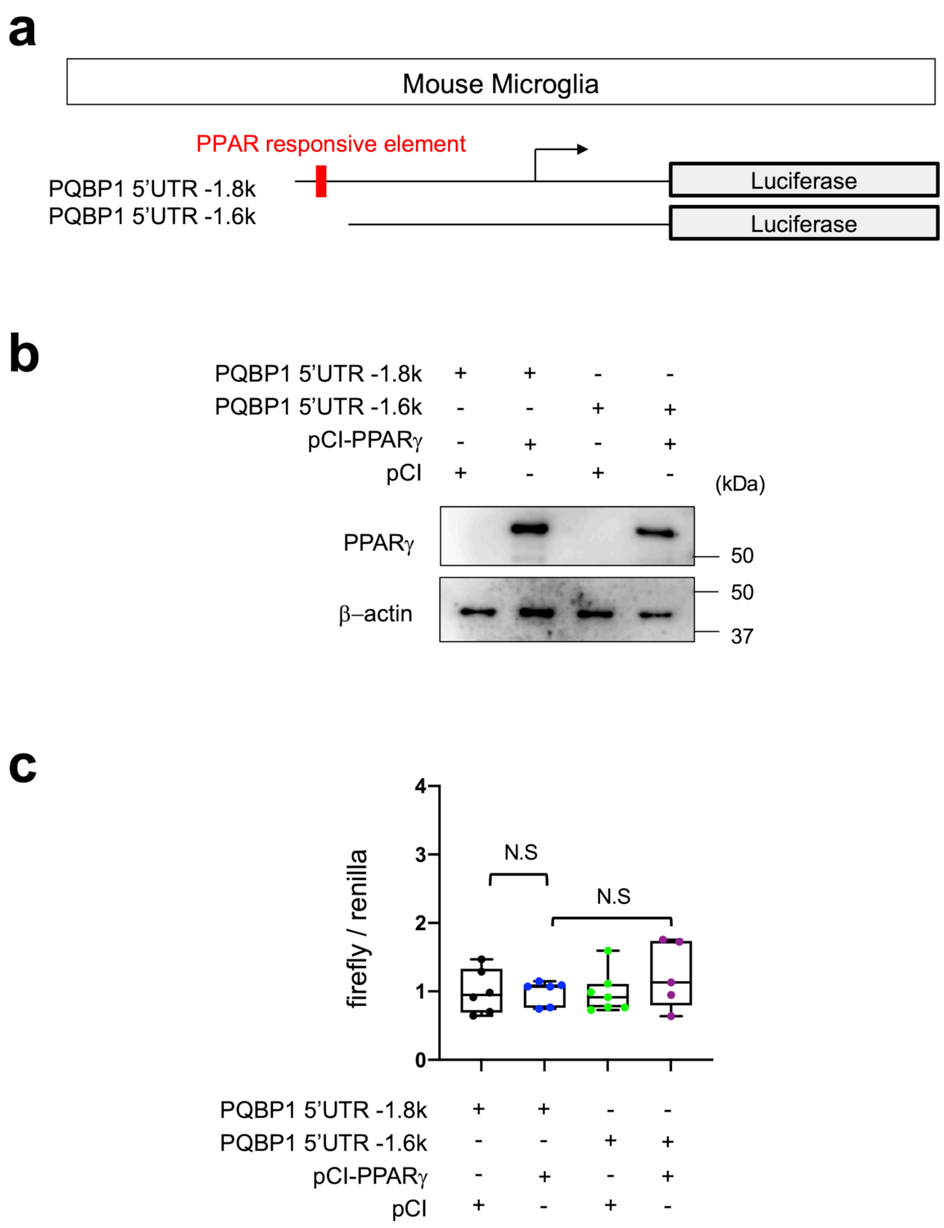


**Supplementary Figure 3**
a) Reporter vectors possessing 1.8 kb or 1.6 kb of the upstream region of mouse *PQBP1* gene were used for luciferase assay in primary microglia.
b) Expression levels of PPARγ in transfected microglia were confirmed by western blot.
c) Luciferase activities were quantified with internal control by Renilla Luciferase Assay system.

#### Supplementary Figure 4

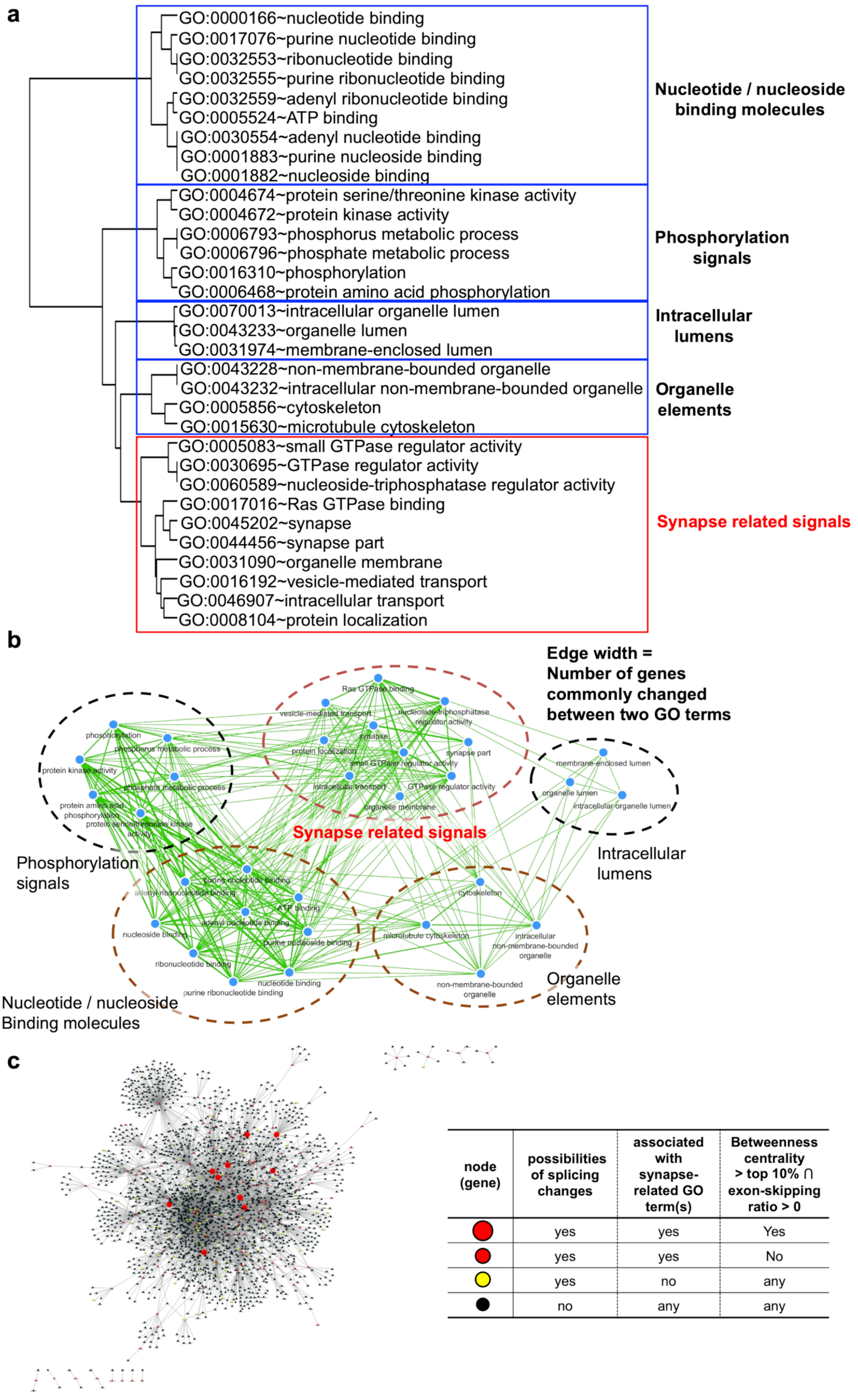


| node (gene) | possibilities of splicing changes | associated with synapse-related GO term(s) | Betweenness centrality > top 10% ∩ exon-skipping ratio > 0 |
|---|---|---|---|
| ● | yes | yes | Yes |
| ● | yes | yes | No |
| ● | yes | no | any |
| ● | no | any | any |

**Supplementary Figure 4**
**Molecular network analysis of AS genes commonly changed in HFD-1W and PQBP1-cKO mice**
a) GO enrichment analysis of 716 genes (1426 exons) that were commonly changed in HFD-1W mice and Syn-cKO mice. GO enrichment revealed five functional groups including the “synapse related signals” group.
b) GO term-based relationship of the five functional groups. Edge width (thickness of green lines) reflects number of genes commonly change between two GO term groups.
c) PPI-based molecular network in which nodes (genes) were categorized as shown in the right table.

#### Supplementary Figure 5

a

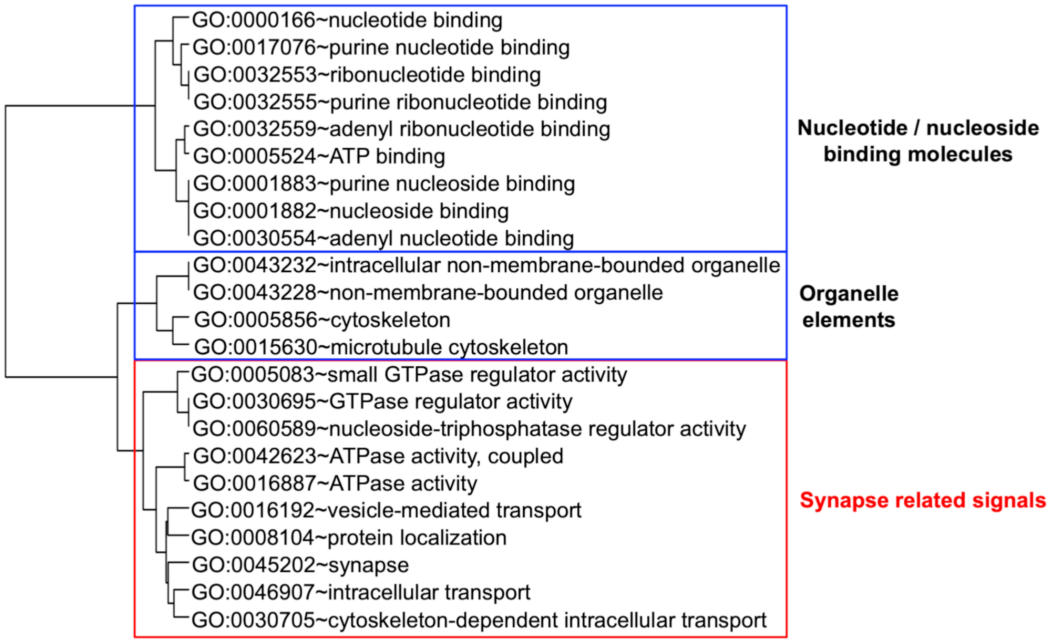


b

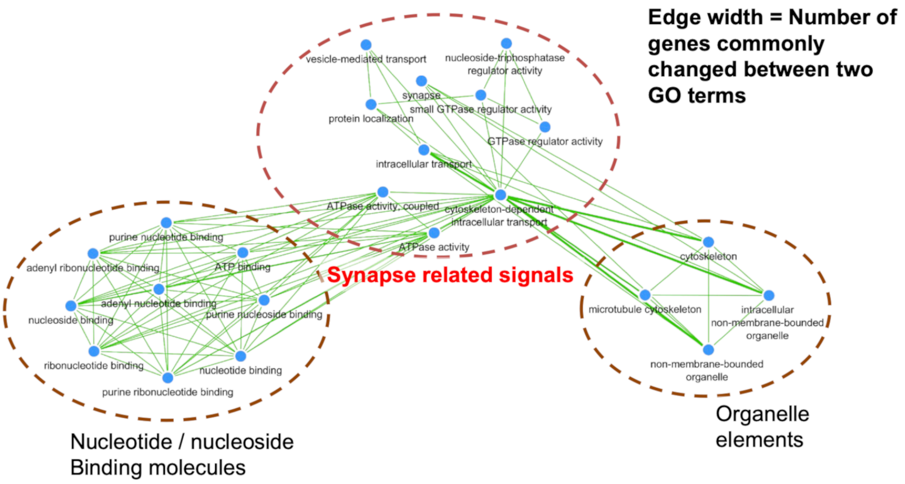


c

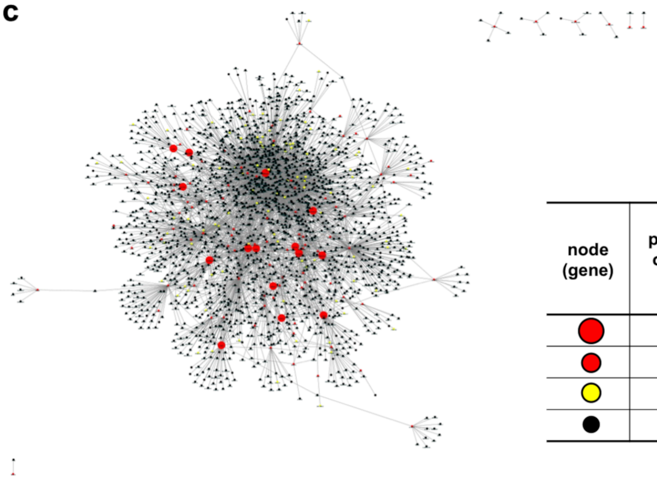

| node (gene) | possibilities of splicing changes | associated with synapse-related GO term(s) | Betweenness centrality > top 10% ∩ exon-skipping ratio > 0 |
|---|---|---|---|
| large red circle | yes | yes | Yes |
| red circle | yes | yes | No |
| yellow circle | yes | no | any |
| black circle | no | any | any |

**Supplementary Figure 5**
**Molecular network analysis of AS genes commonly changed in HFD-6W and PQBP1-cKO mice**
a) GO enrichment analysis of 695 genes (1315 exons) that were commonly changed in HFD-6W mice and Syn-cKO mice. GO enrichment revealed three functional groups including the “synapse related signals” group.
b) GO term-based relationship of the three functional groups. Edge width (thickness of green lines) reflects number of genes commonly change between two GO term groups.
c) PPI-based molecular network in which nodes (genes) were categorized as shown in the right table.

#### Supplementary Figure 6

a

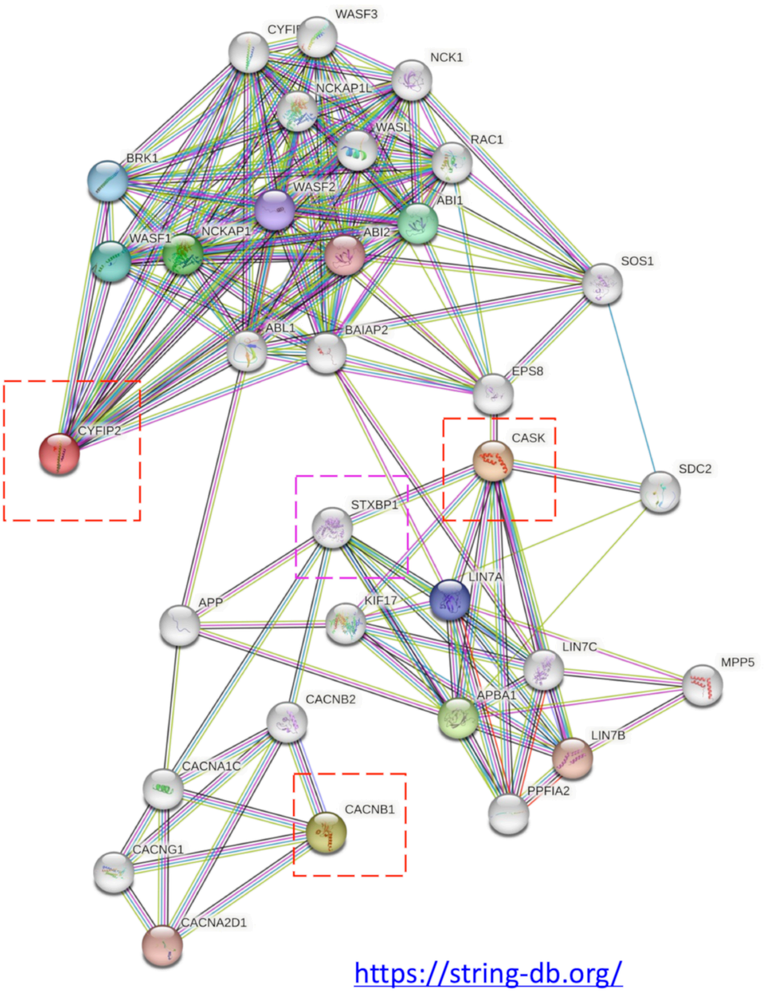


b

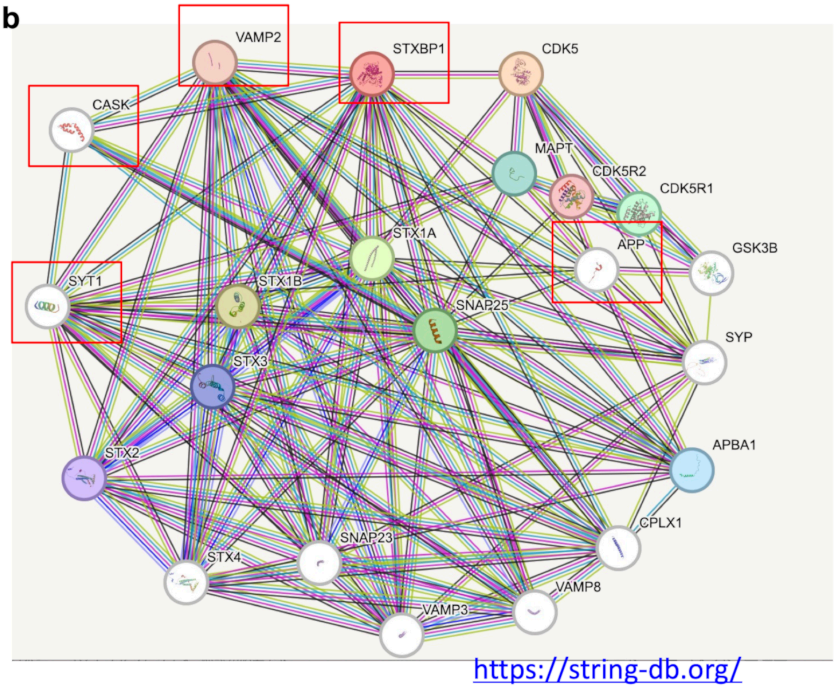


**Supplementary Figure 6**
**Core network composed of key AS genes**
a) Molecular network including three key genes is generated by String ver. 11.5 (https://string-db.org/). The key genes (Cyfip2, Cask and Cacnb1) were indicated by squares of red dot line. STXBP1 was squared by magenta dot line. It is of note that amyloid precursor protein (APP) (blue dot line) connects three groups around each key gene.
b) Another search of molecular network around Cask and Syt1 by String ver. 11.5 (https://string-db.org/).

#### Supplementary Figure 7

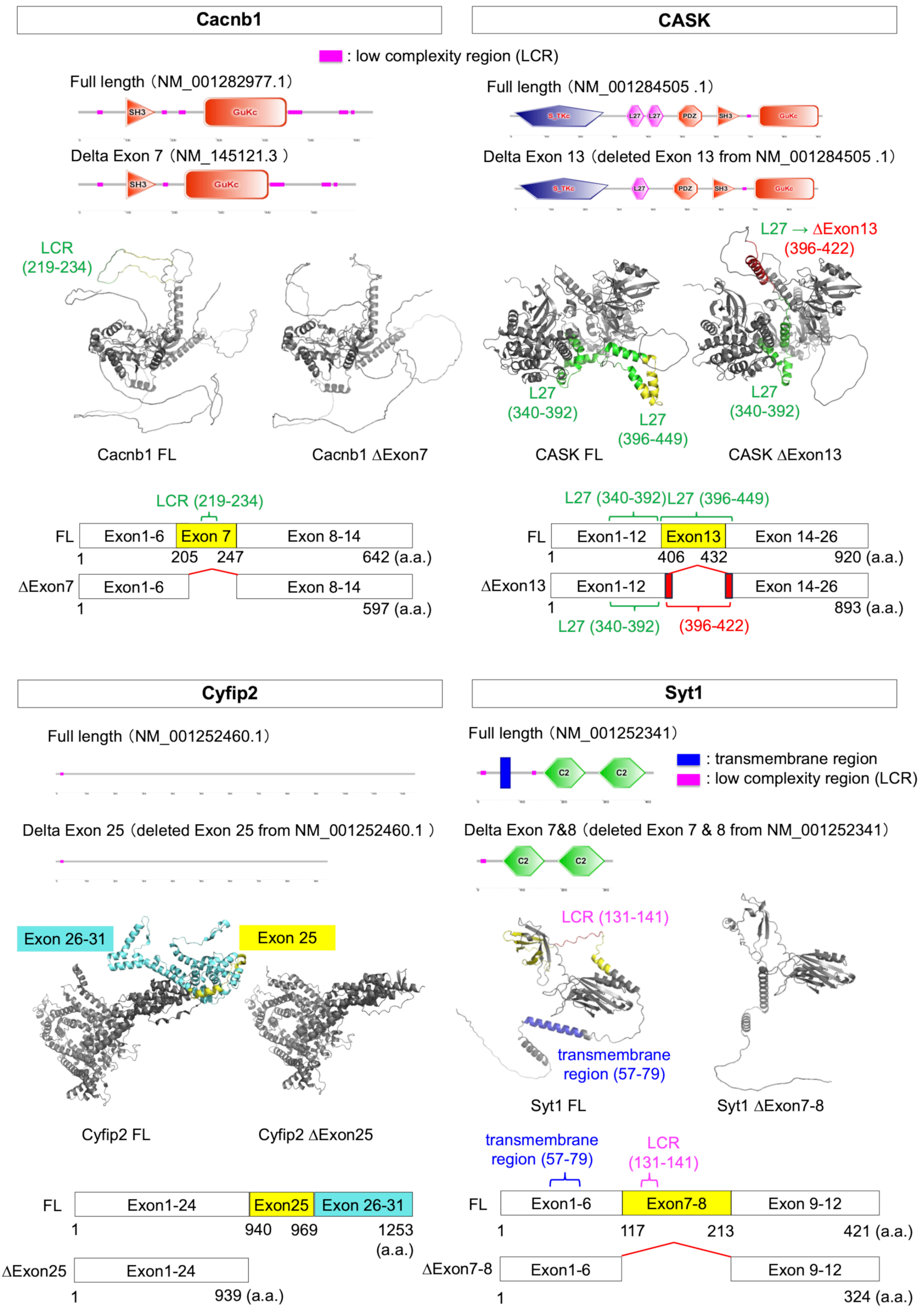


**Supplementary Figure 7**
**Molecular structure of AS isoforms of the HFD-PQBP1 axis target proteins**
In each proteins, upper panels show domain structure of full-length and HFD-induced AS isoforms of target proteins; **m**iddle panels show 3D structures of the full-length and HFD-induced AS isoforms; lower panels show skipped exon positions in protein structures.

### Supplementary Figure 8

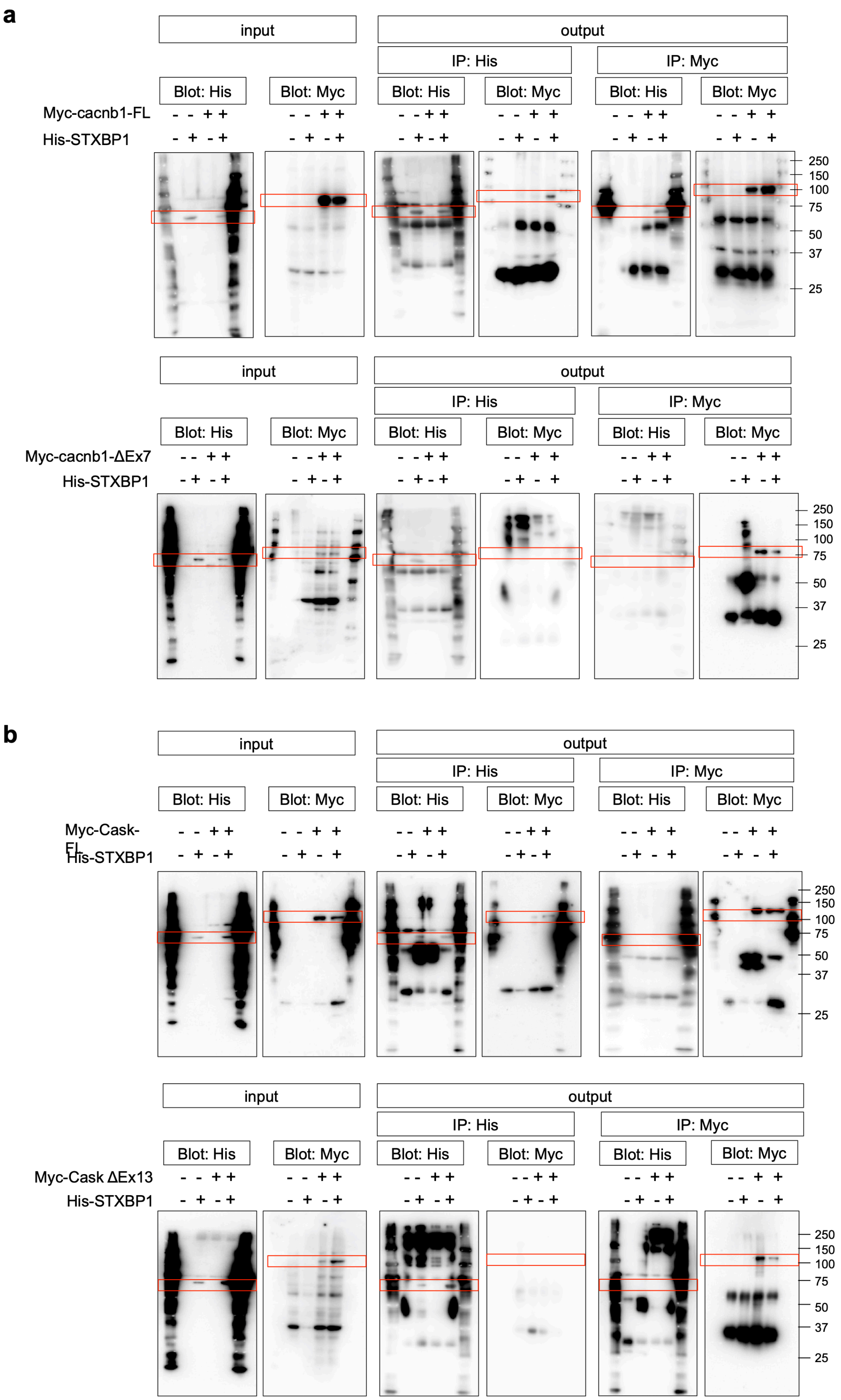


**Supplementary Figure 8**
**Interaction between STXBP1 and Cacnb1 or Cask**

a) Interaction of STXBP1 with full-length and HFD-induced AS isoform of Cacnb1 proteins were analyzed by immunoprecipitation.

b) Interaction of STXBP1 with full-length and HFD-induced AS isoform of Cask proteins were analyzed by immunoprecipitation.

### Supplementary Figure 9

**a**

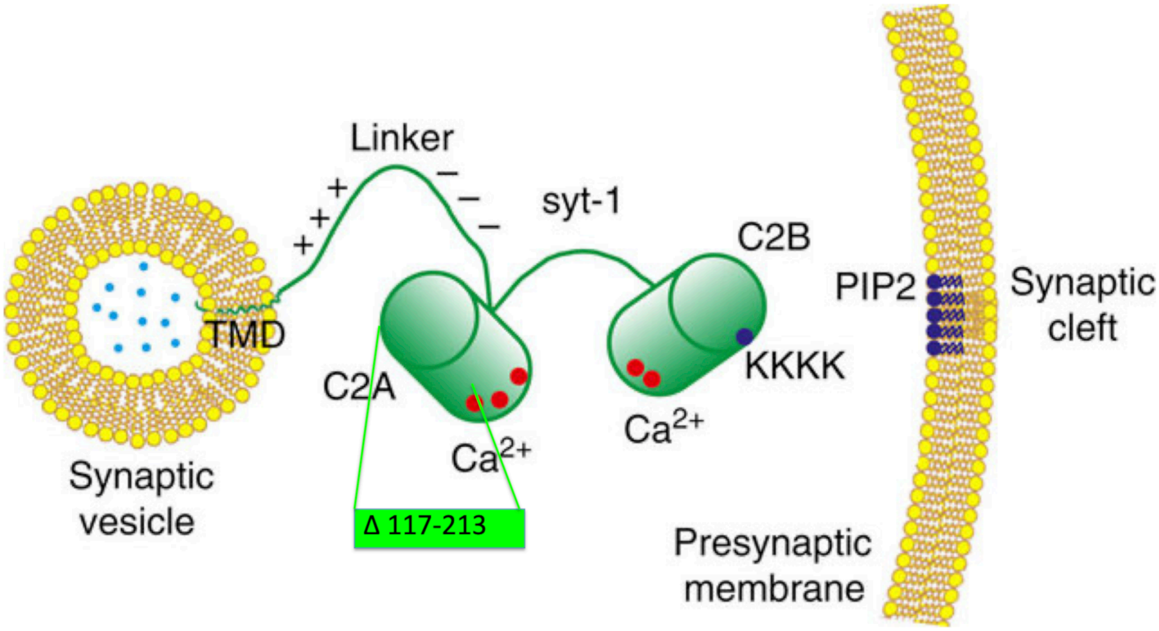


Modified from Lin, CC., Seikowski, J., Pérez-Lara, A. et al. Control of membrane gaps by synaptotagmin-Ca2+ measured with a novel membrane distance ruler. Nat Commun 5, 5859 (2014). https://doi.org/10.1038/ncomms6859

**b**

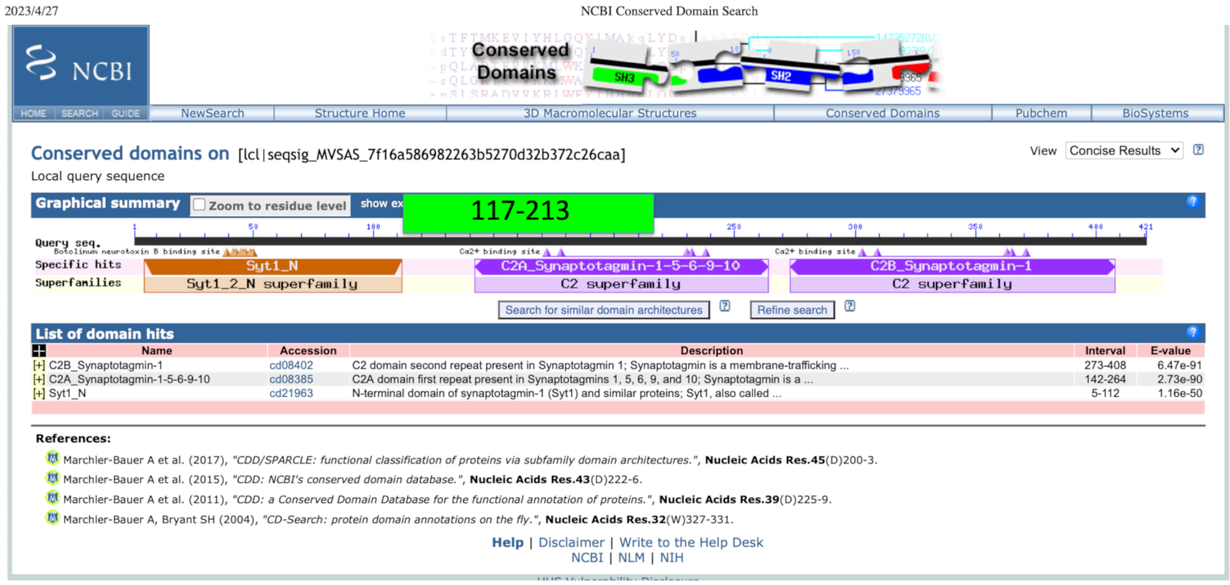

2023/4/27 NCBI Conserved Domain Search

NCBI — HOME | SEARCH | GUIDE | NewSearch | Structure Home | 3D Macromolecular Structures | Conserved Domains | Pubchem | BioSystems

**Conserved domains on** [lcl|seqsig_MVSAS_7f16a586982263b5270d32b372c26caa]

Local query sequence

View Concise Results

**Graphical summary** Zoom to residue level show ex



Search for similar domain architectures | Refine search

**List of domain hits**

| Name | Accession | Description | Interval | E-value |
|---|---|---|---|---|
| [+] C2B_Synaptotagmin-1 | cd08402 | C2 domain second repeat present in Synaptotagmin 1; Synaptotagmin is a membrane-trafficking ... | 273-408 | 6.47e-91 |
| [+] C2A_Synaptotagmin-1-5-6-9-10 | cd08385 | C2A domain first repeat present in Synaptotagmins 1, 5, 6, 9, and 10; Synaptotagmin is a ... | 142-264 | 2.73e-90 |
| [+] Syt1_N | cd21963 | N-terminal domain of synaptotagmin-1 (Syt1) and similar proteins; Syt1, also called ... | 5-112 | 1.16e-50 |


**Help** | Disclaimer | Write to the Help Desk
NCBI | NLM | NIH

**Supplementary Figure 9**
**HFD-induced AS and domain structure of Syt1**
a) Exon 7 & 8 of Syt1 skipped by HFD corresponds to C2A domain.
b) Mapping exon 7 & 8 regions to domain structure predicted by NCBI conserved domain search.

#### Supplementary Figure 10

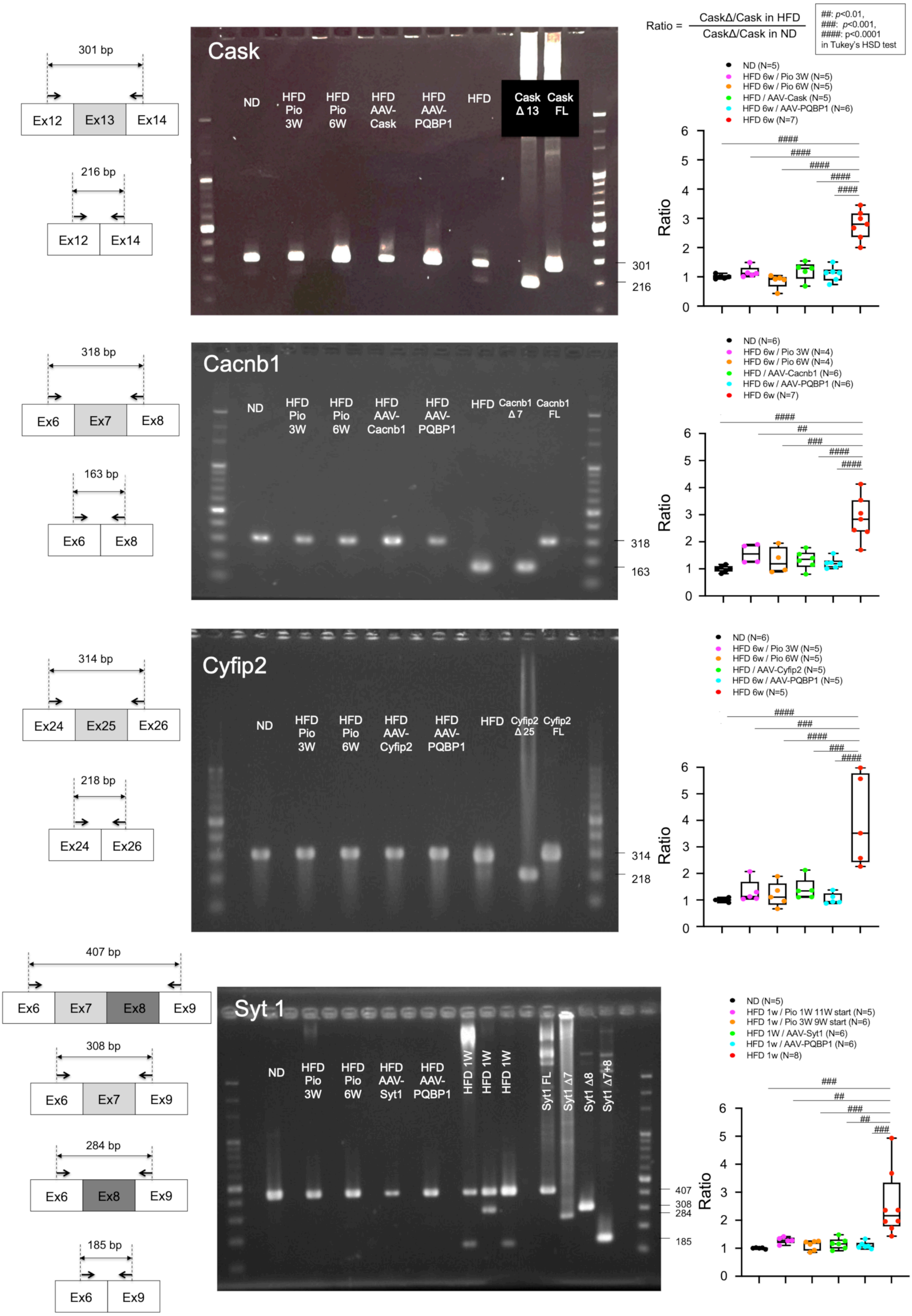


**Supplementary Figure 10**

**HFD-induced AS isoforms of target genes detected by RT-PCR with cerebral cortex tissues**

RT-PCR was performed with cerebral cortex tissues prepared from normal diet-fed and high fat diet-fed mice with or without treatment (pioglitazone and AAV-PQBP1).

#### Supplementary Figure 11

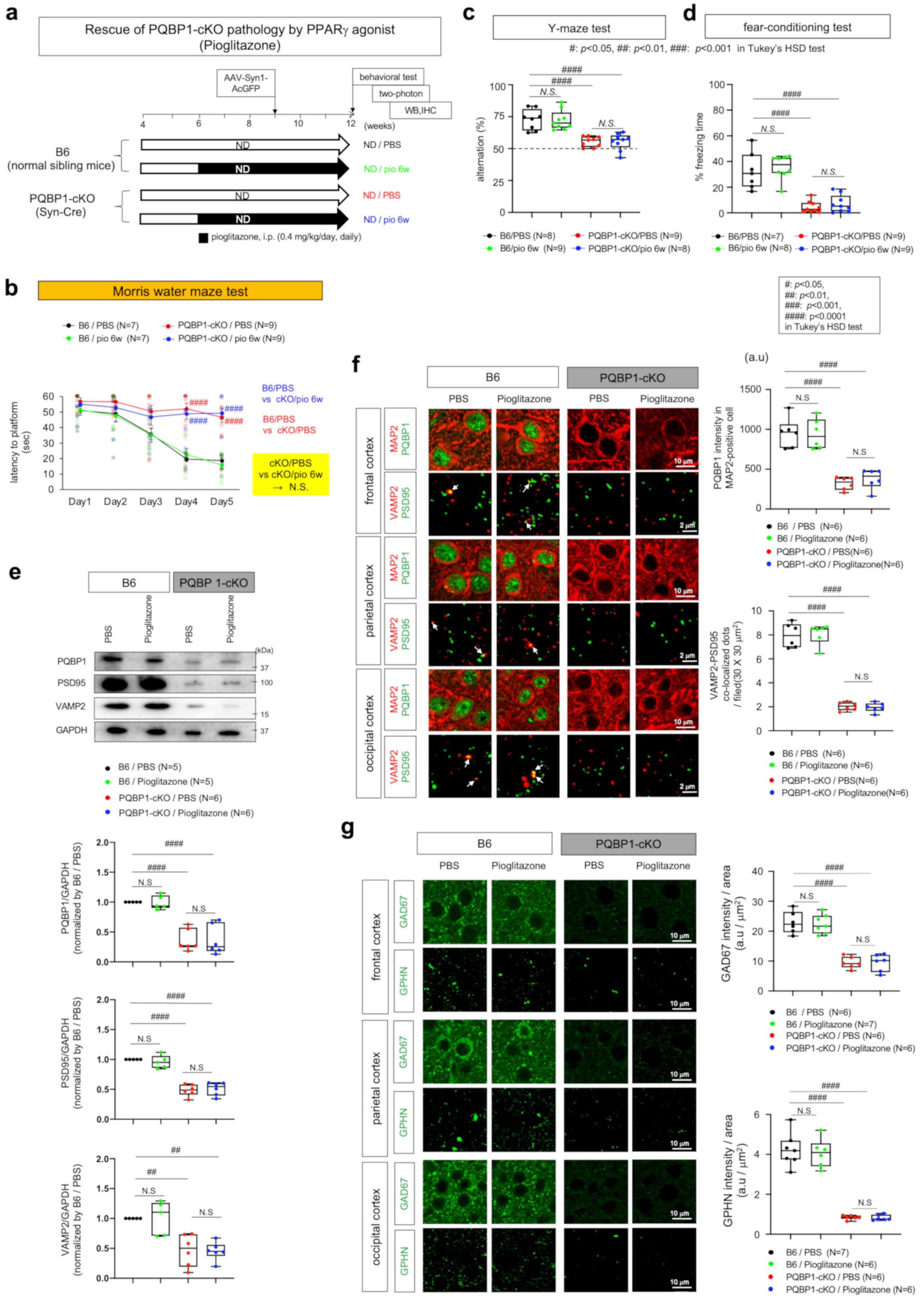


**Supplementary Figure 11**
**Rescue of PQBP1-cKO spine pathology by PPARg agonist**
a) Protocol of the rescue experiment for synapse pathology of Syn-cKO mice by pioglitazone. 4 mouse groups were prepared.
b) Results of Morris water maze-test in the 4 mouse groups.

c) Results of Y maze-test in the 4 mouse groups.
d) Results of fear conditioning test in the 4 mouse groups.
e) Western blot analysis of PQBP1, PSD95 (post-synapse marker) and VAMP2 (pre-synapse marker) in total cerebral cortex tissues of the 4 mouse groups. Right graphs show quantitative analyses of the results.
f) Immunohistochemistry of PQBP1 in MAP2-positive neurons and of VAMP2-PSD95 co-staining for mature synapses in 4 mouse groups. Right graphs show quantitative analyses of signal intensities of neuronal PQBP1 and of numbers of mature synapses. The mean value of three cortex areas was used as a representative value for a mouse.
g) Immunohistochemistry of GAD67 (pre-synapse marker of inhibitory synapses) and GPHN (post-synapse marker of inhibitory synapses).

**Supplementary Figure 12**

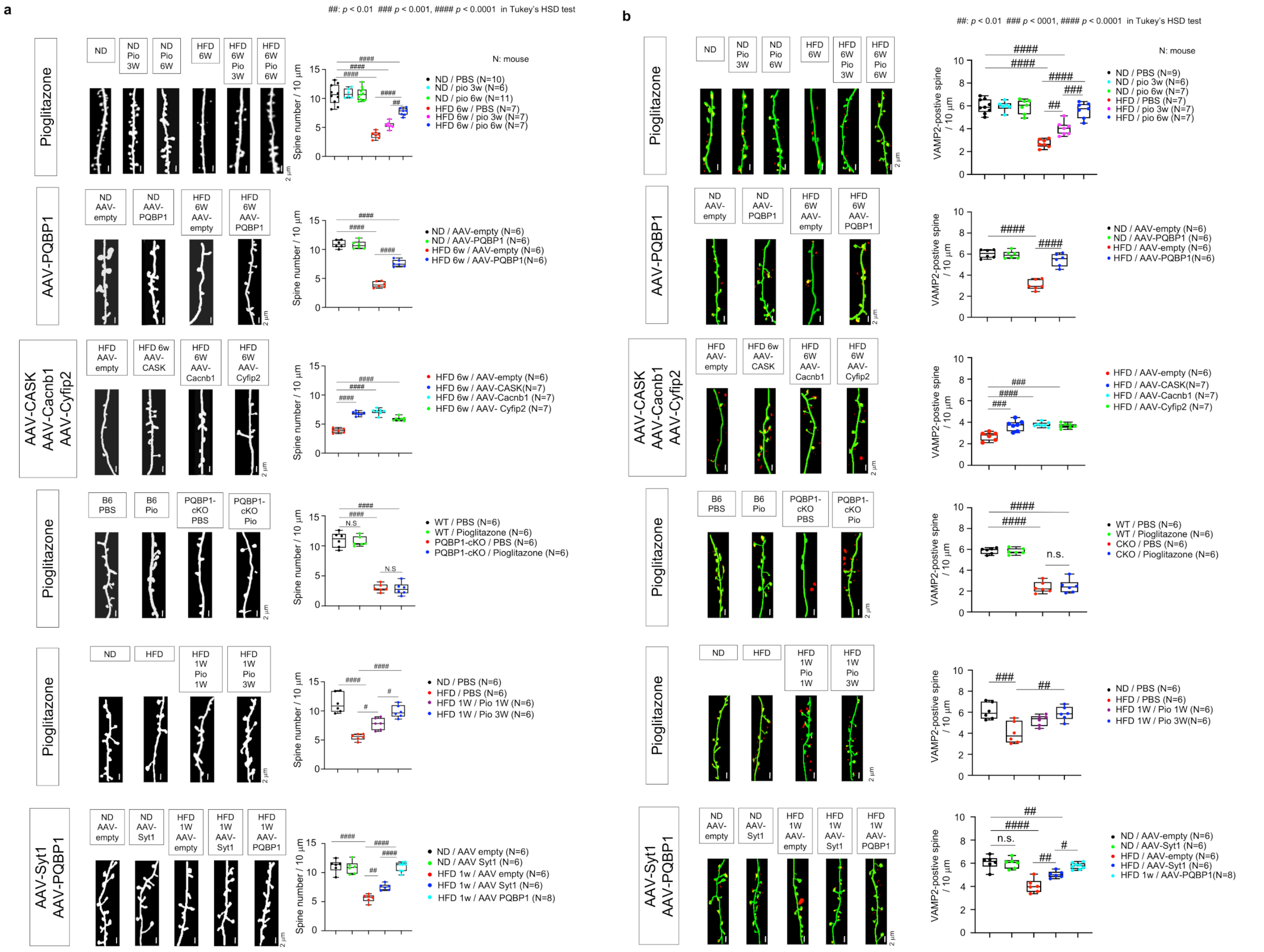


#### Supplementary Figure 12
#### Two-photon microscopic analyses of dendritic spines in rescue experiments

a) Dendritic spines were analyzed by AAV-EGFP with synapsin 1 promoter.

b) VAMP2-positive dendritic spines were analyzed by AAV-EGFP with synapsin 1 promoter and AAV-VAMP2-mCherry.

### Supplementary Figure 13

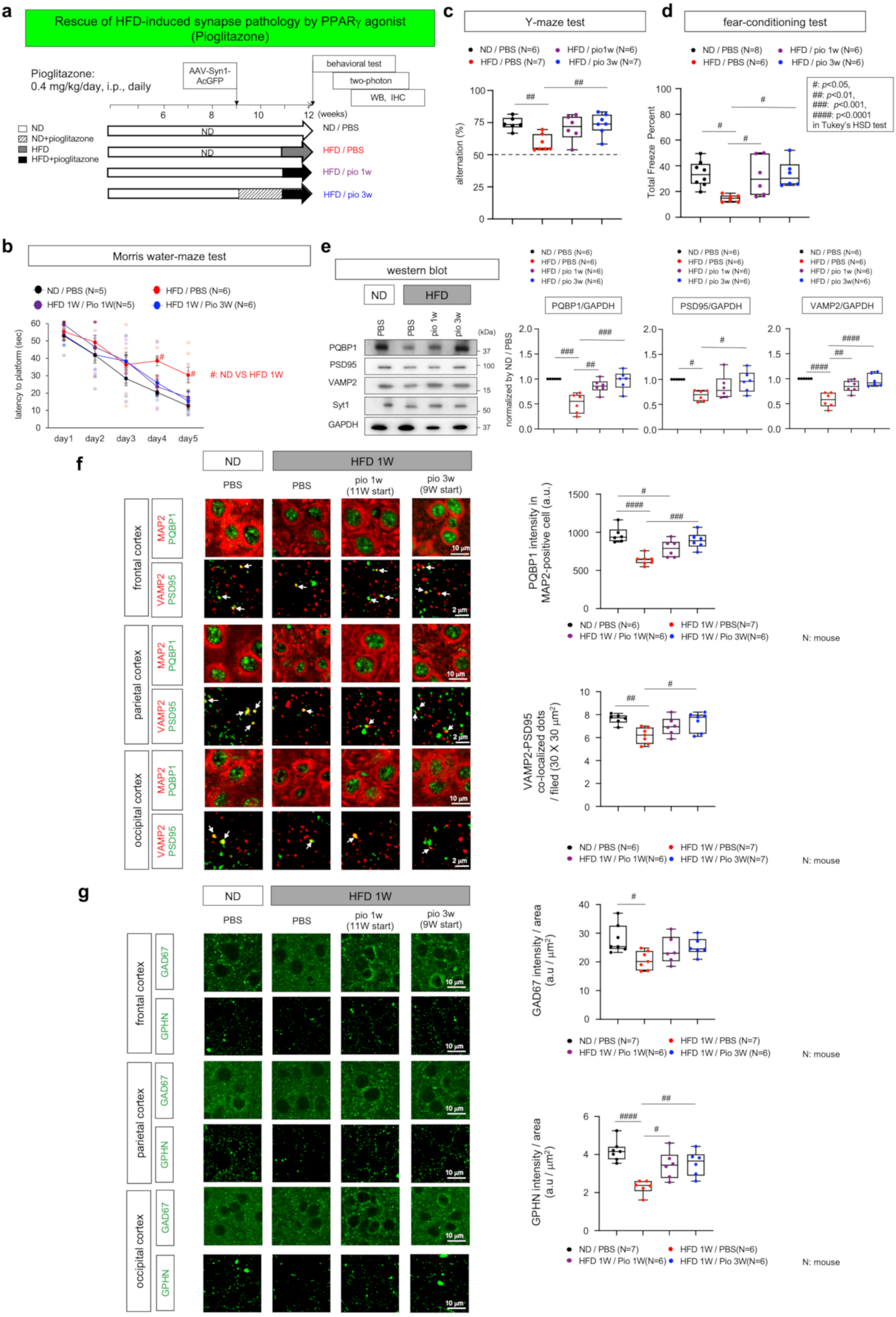

**Supplementary Figure 13**
**Rescue of HFD1W-induced synapse pathology by PPARγ agonist**
a) Protocol of the rescue experiment for synapse pathology of HFD1W mice by pioglitazone. Four mouse groups were prepared.
b) Results of Morris water maze-test in the 4 mouse groups.
c) Results of Y maze-test in the 4 mouse groups.
d) Results of fear conditioning test in the 4 mouse groups.
e) Western blot analysis of PQBP1, PSD95 (post-synapse marker) and VAMP2 (pre-synapse marker) in total cerebral cortex tissues of the 4 mouse groups. Right graphs show quantitative analyses of the results.
f) Immunohistochemistry of PQBP1 in MAP2-positive neurons and of VAMP2-PSD95 co-staining for mature synapses in 4 mouse groups. Right graphs show quantitative analyses of signal intensities of neuronal PQBP1 and of numbers of mature synapses. The mean value of three cortex areas was used as a representative value for a mouse.
g) Immunohistochemistry of GAD67 (pre-synapse marker of inhibitory synapses) and GPHN (post-synapse marker of inhibitory synapses).

### Supplementary Figure 14

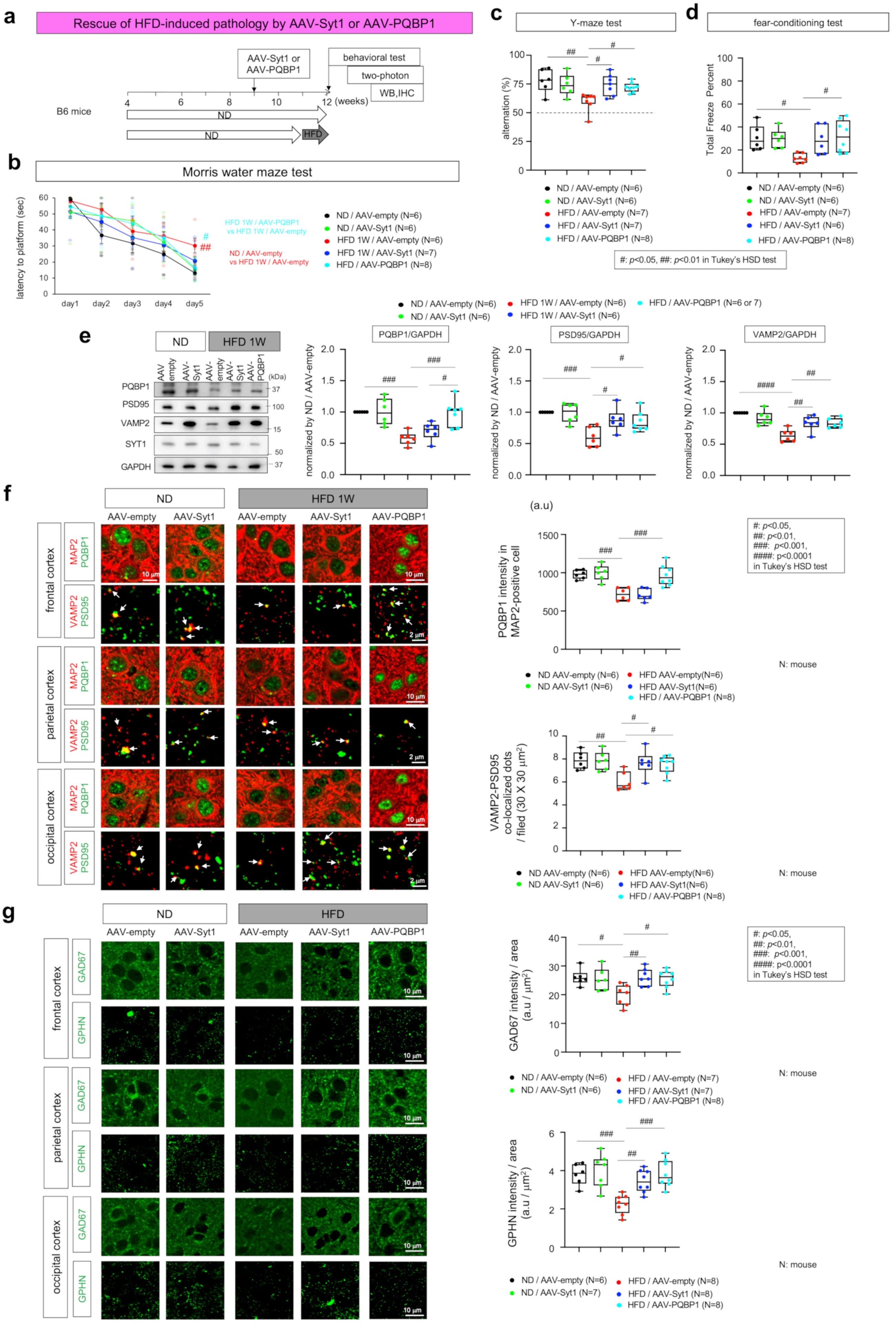

**Supplementary Figure 14**
**Rescue of HFD1W-induced pathology by AAV-Syt1 or AAV-PQBP1**

a) Protocol of the rescue experiment for synapse pathology of HFD1W mice by AAV-PQBP1. Two mouse groups were prepared.
b) Results of Morris water maze-test in the 4 mouse groups.
c) Results of Y maze-test in the 4 mouse groups.
d) Results of fear conditioning test in the 4 mouse groups.
e) Western blot analysis of PQBP1, PSD95 (post-synapse marker) and VAMP2 (pre-synapse marker) in total cerebral cortex tissues of the 4 mouse groups. Right graphs show quantitative analyses of the results.
f) Immunohistochemistry of PQBP1 in MAP2-positive neurons and of VAMP2-PSD95 co-staining for mature synapses in 4 mouse groups. Right graphs show quantitative analyses of signal intensities of neuronal PQBP1 and of numbers of mature synapses. The mean value of three cortex areas was used as a representative value for a mouse.
g) Immunohistochemistry of GAD67 (pre-synapse marker of inhibitory synapses) and GPHN (post-synapse marker of inhibitory synapses).

**Supplementary Table 1**
**AS-changed exons in WT-HFD1W**

**Supplementary Table 2**
**AS-changed exons in WT-HFD6W**

**Supplementary Table 3**
**AS-changed exons in Syn-cKO**

**Supplementary Table 4**
**commonly AS-changed exons between WT-HFD1W and Syn-cKO**

**Supplementary Table 5**
**commonly AS-changed exons between WT-HFD6W and Syn-cKO**

**Supplementary Table 6**
**GO terms enriched by commonly AS-changed exons between WT-HFD1W and Syn-cKO**

**Supplementary Table 7**
**GO terms enriched by commonly AS-changed exons between WT-HFD6W and Syn-cKO**

**Supplementary Table 8**
**Betweenness centrality scores of commonly AS-changed nodes between WT-HFD1W and Syn-cKO**

**Supplementary Table 9**
**Betweenness centrality scores of commonly AS-changed nodes between WT-HFD6W and Syn-cKO**
